# Opposing regulation of short-term memory by basal ganglia direct and indirect pathways that are coactive during behavior

**DOI:** 10.1101/2021.12.15.472735

**Authors:** Yingjun Tang, Hongjiang Yang, Xia Chen, Zhouzhou Zhang, Xiao Yao, Xinxin Yin, Zengcai V. Guo

**Affiliations:** Tsinghua-Peking Joint Center for Life Sciences, Beijing, China, 100084; School of Medicine, Tsinghua University, Beijing, China, 100084; IDG/McGovern Institute for Brain Research, Tsinghua University, Beijing, China, 100084; School of Life Sciences, Tsinghua University, 100084, Beijing, China; Peking-Tsinghua Joint Center for Life Sciences, Beijing, China, 100084

**Author notes:** These authors contributed equally to this work (Yingjun Tang, Hongjiang Yang).

## Abstract

The basal ganglia direct and indirect pathways are viewed to mediate opposing functions in movement. However, this classic model is challenged by recent findings that both pathways are coactive during behavior. We examined the roles of direct (dSPNs) and indirect (iSPNs) pathway spiny projection neurons in a decision-making task with a short-term memory (STM) component. Optogenetic stimulation of cortical-input-defined dSPNs and iSPNs during STM oppositely biased upcoming licking choice, without affecting licking execution. Optogenetically identified dSPNs and iSPNs showed similar response patterns, although with quantitative difference in spatiotemporal organization. To understand how coactive dSPNs and iSPNs play opposing roles, we recorded population activity in frontal cortex and the basal ganglia output nucleus SNr. Stimulation of dSPNs and iSPNs bidirectionally regulated cortical decision variable through the differential modulation of SNr ramping activity. These results reconcile different views by demonstrating that coactive dSPNs and iSPNs precisely shape cortical activity in a push-pull balance.

## Introduction

The basal ganglia play crucial roles in diverse functions including motor control and reward-based learning (Albin et al., 1989; Cox and Witten, 2019; DeLong, 1990; Graybiel, 2008; Hikosaka et al., 2000). Dysfunction of the circuit causes severe movement disorders like Parkinson’s and Huntington’s diseases (Albin et al., 1989; DeLong, 1990; Graybiel, 2008), as well as cognitive deficits such as working memory and attention (Aarsland et al., 2017; Simpson et al., 2010). In the classic view that is often referred to as the ‘go/no-go’ model (Albin et al., 1989; DeLong, 1990), the basal ganglia direct and indirect pathways function to facilitate and suppress motor behaviors, respectively. The direct pathway facilitates movement by inhibiting the output nuclei, the substantial nigra pars reticulata (SNr) and the internal segment of the globus pallidus (GPi), which in turn inhibit an array of thalamic and brainstem nuclei by their GABAergic projections. The indirect pathway, which projects to the external segment of the globus pallidus (GPe) and subthalamic nucleus (STN), suppresses movement by disinhibiting the output nuclei of the basal ganglia. The ‘go/no-go’ model is powerful in explaining symptoms caused by the Parkinson’s and Huntington’s diseases. The striatal spiny projection neurons (SPNs) of the direct pathway primarily express the D1 dopamine receptor, while those of the indirect pathway mostly express the D2 dopamine receptor, enabling pathway-specific perturbations (Gerfen et al., 1990). Perturbation of dSPNs and iSPNs promotes and demotes movements including locomotion and level pressing (Bateup et al., 2010; Chen et al., 2021; Kravitz et al., 2010; Oldenburg and Sabatini, 2015), although the effect of perturbation can be asymmetric or behavioral-epoch dependent (Lee and Sabatini, 2021; Oldenburg and Sabatini, 2015; Tecuapetla et al., 2016). The simplest form of the ‘go/no-go’ model suggests that dSPNs and iSPNs would not be co-active during a single well-defined movement because the two pathways function antagonistically.

However, accumulating evidence has repeatedly demonstrated that neurons in the two pathways are co-activated during movement initiation and termination (Barbera et al., 2016; Chen et al., 2021; Cui et al., 2013; Jin et al., 2014; Markowitz et al., 2018). For example, population recording of calcium activity in dSPNs and iSPNs indicates that both pathways are active when mice initiate movements to shuttle between food magazine and levers (Cui et al., 2013). Single neuronal recording of calcium activity demonstrates that dSPNs and iSPNs are concurrently active during locomotion initiation and termination (Barbera et al., 2016), although activity in the direct and indirect pathways also shows decorrelation at sub-second timescales (Chen et al., 2021; Markowitz et al., 2018; Sippy et al., 2015). Electrical recording of optically identified dSPNs and iSPNs provides higher temporal resolution and further demonstrates that both types of neurons are active during level pressing (Jin et al., 2014). Although dSPNs and iSPNs exhibit similar pattern of activation, there also exists quantitative difference in response especially during movement transitions or to reward (Chen et al., 2021; Jin et al., 2014; Markowitz et al., 2018; Shin et al., 2018; Sippy et al., 2015).

Concurrent activation of the direct and indirect pathway projection neurons theoretically could be related to the converging cortical inputs and the dynamic activation during different phases of movement. Striatal neurons receive convergent inputs from multiple cortical regions (Wall et al., 2013). Although different zones of the striatum receive coarse topographical projections from the cortex, there are significant overlaps in projections between adjacent zones (Hintiryan et al., 2016; Hunnicutt et al., 2016). Distinct cortical input targeting the same region of striatum may affect behavior in opposing ways (Lee et al., 2019). Movement also has complicated dynamics involving changing of movement velocity and direction through acceleration and deceleration. Accordingly, motor cortex neurons exhibit dynamic modulation that could drive dSPNs and iSPNs with complicated firing patterns (Churchland et al., 2012). With these considerations, we aim to study how cortex-input-defined dSPNs and iSPNs function in a pseudo-steady state (i.e. the delay period) of a tactile-based decision task (Guo, Li, et al. 2014).

During the delay period in the decision task, the mouse anterior lateral motor cortex (ALM) plays a causal role for short-term memory (STM) (Guo, Li, et al. 2014). A large fraction of ALM neurons exhibit trial-type specific activity that predicts upcoming licking directions (Li et al., 2016). We thus focused on striatal dSPNs and iSPNs that were transsynaptically labelled from ALM. By employing input-defined and pathway-specific labelling, optogenetic stimulation and recording, we found that dSPNs and iSPNs had opposing roles in regulating STM, consistent with the classic view, while they had similar response patterns in the behavioral task, consistent with recent findings of concurrent activation. Reconciling different views requires recording and perturbing dSPNs and iSPNs in the same behavioral task and simultaneously monitoring downstream activity to understand the effects of perturbation. We found that concurrent activity in dSPNs and iSPNs bidirectionally shifted ALM population activity underlying STM to oppositely bias upcoming choice. And the opposing effects was through their bidirectional modulation of non-selective activity in SNr. These results underscore the importance of using input-defined cell-type specific perturbation and recording to study the basal ganglia direct and indirect pathways.

## Results

### Stimulation of cortical-input-defined direct and indirect pathway SPNs oppositely regulates STM

Previous studies have demonstrated that the direct and indirect pathways mediate opposing roles in movement (Chen et al., 2021; Kravitz et al., 2010; Lee and Sabatini, 2021; Oldenburg and Sabatini, 2015; Tecuapetla et al., 2016). However, it remains unknown if these pathways function antagonistically in STM. To address this, we employed input-defined labelling with transgenic Cre driver lines targeting dSPNs and iSPNs. We injected serotype 1 virus (scAAV1-hSyn-flex-EGFP) in ALM in D1-Cre or D2-Cre mice (Figure 1A and C). AAV1 has been shown to specifically transduce postsynaptically connected neurons in a synaptic transmission dependent way (Zingg et al., 2017; Zingg et al., 2020). Consistently, we observed widespread labelling of dSPNs and iSPNs in ALM projection zone (Figure 1 B and D; Figure S1). ALM-input defined dSPNs (dSPNALM) projected extensively in SNr while ALM-input defined iSPNs (iSPNALM) mainly projected to GPe, with little cross-projections that is consistent with known anatomical connections (Albin et al., 1989; Gerfen et al., 1990; Kawaguchi et al., 1990). Apart from ipsilateral projections, dSPNALM and iSPNALM also sent axons crossing corpus callosum to reach contralateral targets. The projection pattern in the contralateral hemisphere was similar, although the intensity of projection was lower (Figure S1).

**Fig. 1.**
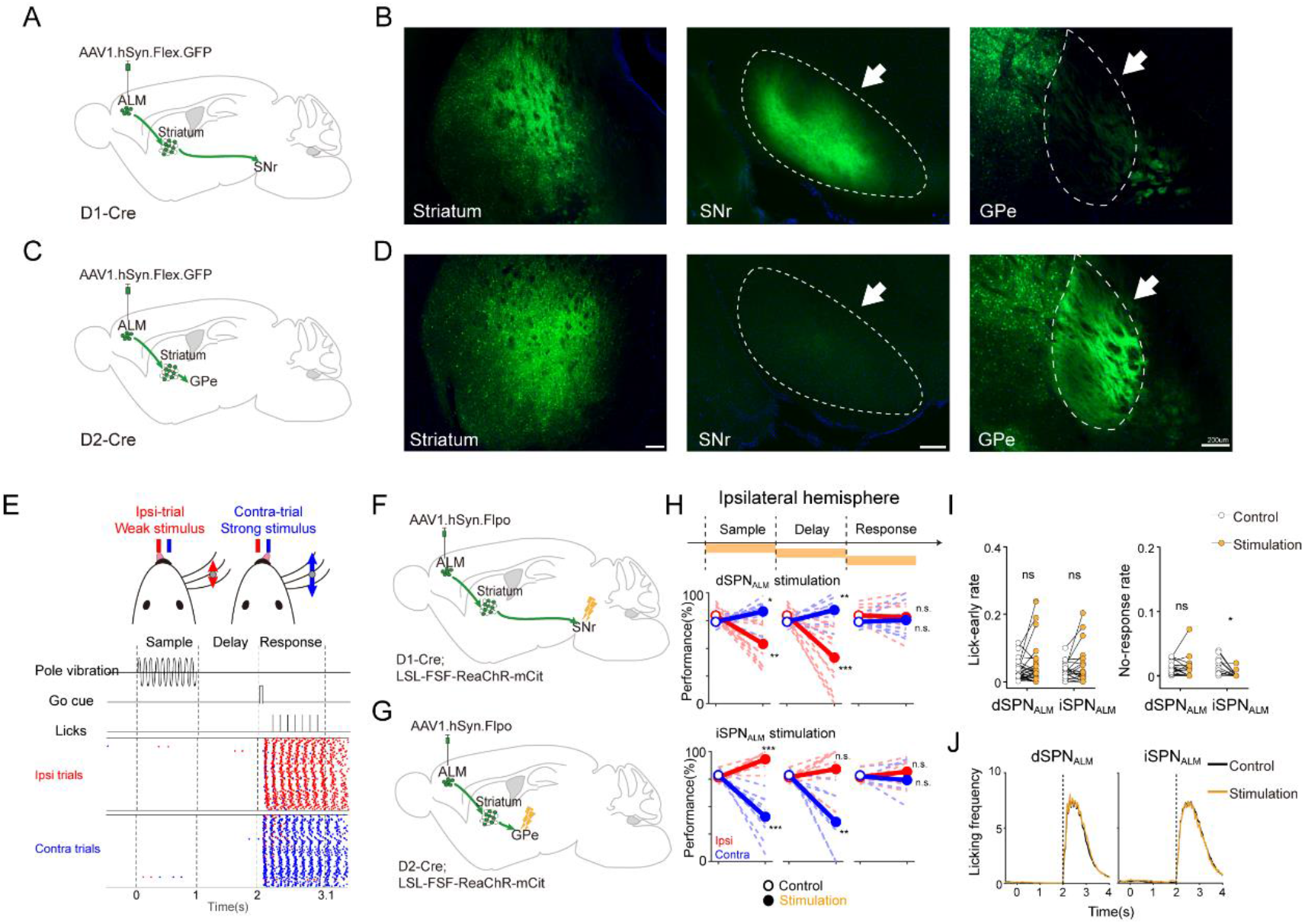
Stimulation of ALM-input defined direct and indirect pathway SPNs demonstrates their opposing roles in STM. (A) Schematic of labelling of ALM-input defined direct pathway SPNs (dSPN_ALM_) by injecting anterograde transsynaptic virus in ALM of D1-Cre mice. (B) dSPN_ALM_ neurons are widely distributed in the striatum (left) and their axons project to SNr (middle) but not to GPe (right). (C) Schematic of labelling of ALM-input defined indirect pathway SPNs (iSPN_ALM_) (D) iSPN_ALM_ neurons are widely distributed in the striatum (left) and their axons mainly target GPe (right) but not SNr (middle). (E) The tactile-based decision task with a STM component. Top: Head-fixed mice were trained to lick the ipsilateral or contralateral water spout based on the strength of whisker stimulation. Contra and ipsi denote the side relative to the optogenetically stimulated left hemisphere. Middle: behavioral events (pole vibration, go signal and licking during the sample, delay and response epochs). Bottom: an example behavioral session demonstrates that mice can withhold licking during the sample and delay epochs. (F) Schematic of labelling dSPN_ALM_ with red-shifted channelrhodopsin (ReaChR). For activation, an optic fiber was implanted to stimulate the projection axons in SNr. (G) Schematic of labelling iSPN_ALM_ with ReaChR. For activation, an optic fiber was implanted to stimulate the projection axons in GPe. (H) Stimulation of dSPN_ALM_ during the sample or delay epoch produced a contralateral bias while stimulation of iSPN_ALM_ produced an ipsilateral bias. Thin line, individual mice (n = 13 D1-Cre, n= 10 D2-Cre mice). Thick line, mean. (I) Stimulation of dSPN_ALM_ and iSPN_ALM_ did not affect lick-early rate (left). Stimulation of iSPN_ALM_ slightly reduced no-response rate (from 0.77 ± 0.26% to 0.13 ± 0.09%, Mean ± SEM). Thin line, individual behavioral sessions (n = 32 from 13 D1-Cre mice; n= 22 from 10 D2-Cre mice). (J) Stimulation of dSPN_ALM_ and iSPN_ALM_ during response period did not affect licking. Scale bar, 200 μm. *, **, & *** denote paired two-sided t-test with statistical significance P < 0.05, P < 0.01, & P < 0.001, respectively. ns, not significant.

To check whether dSPN_ALM_ and iSPN_ALM_ differentially regulate cognitive function, we adopted a tactile-based decision task with a STM component (Guo et al., 2014b). Mice discriminated the strength of a vibration stimulus using their whiskers and reported the strength with directional licking (ipsilateral or contralateral) to obtain a milk reward (Wang et al., 2021). Whisker stimulation was limited to the sample epoch and a time delay (1.0 s) separated the sensory instruction and the subsequent behavioral response. Mice withheld licking during the sample and delay epoch, and the rate of licking before the response cue was low (Figure 1E, Methods). Thus mice must use STM to link whisker stimulation with correct choice.

To efficiently label dSPN_ALM_ and iSPN_ALM_ for optogenetic stimulation, we injected anterograde transsynaptic virus (AAV1-hSyn-FLPO) in double-restricted transgenic mice (R26 LSL FSF ReaChR-mCitrine x D1-cre or D2-cre, Figure 1F, G). As dSPN_ALM_ and iSPN_ALM_ were widely distributed in the ALM projection zone that spanned ~2-3 millimeters in three dimensions (Figure S1), we implanted optical fibers over SNr and GPe to efficiently and specifically activate striatonigral and striatopallidal projections, respectively (Figure 1F, G). Temporal-specific optogenetic stimulation was deployed during the sample, delay or response epoch in ~25% randomly selected trials. Unilateral stimulation of dSPN_ALM_ axons during the sample or delay epoch significantly increased performance on contra-trials (P < 0.05 in sample epoch, P < 0.01 in delay epoch, t-test) and significantly reduced performance on ipsi-trials (P < 0.01 in sample epoch and P < 0.001 in delay epoch), producing a contralateral bias (Figure 1H). Stimulation of iSPN_ALM_ axons produced an opposite pattern of behavioral deficits (significant increase for sample epoch stimulation on ipsilateral trials, P < 0.001, t-test; significant decrease on contralateral trials, sample epoch, P < 0.001, delay epoch, P < 0.01). As stimulation of pyramidal tract neurons in the ALM output layer causes a contralateral bias (Li et al., 2015), stimulation of dSPN_ALM_ and iSPN_ALM_ presumably promotes and demotes ALM activity respectively to produce a directional bias. Stimulation at the same laser power during the response epoch had little effect on performance, suggesting that the manipulation affected STM rather than motor execution (Figure 1H). Notably, stimulation of the two pathways did not affect lick-early rate and licking rate (Figure 1I, J). Stimulation of the indirect pathway decreased no-response rate but the reduction was small (from 0.77 ± 0.26% to 0.13 ± 0.09%, Mean ± SEM, Figure 1I). These results demonstrate that the direct and indirect pathways oppositely regulate STM. By increasing the stimulation power, we also observed that stimulation of iSPN_ALM_ but not dSPN_ALM_ axons induced a significant increase of lick-early rate (Figure S2A, B), indicating that the role of direct and indirect pathways can be complex under strong stimulation.

Anterograde virus injected in the left ALM also labelled dSPN_ALM_ and iSPN_ALM_ in the right striatum (Figure S3). As dSPN_ALM_ and iSPN_ALM_ in the left and right striatum both received input from the left ALM, stimulation in the left and right hemispheres could cause a similar behavioral bias (i.e. a bias from left- to right-direction for the stimulation of direct pathway and a bias from right- to left-direction for stimulation of the indirect pathway). Alternatively, stimulation in the left and right hemisphere could cause an opposite directional bias (i.e. similar directional bias relative to the stimulated side). To differentiate the two possibilities, we stimulated dSPN_ALM_ and iSPN_ALM_ in the right hemisphere (Figure S3A). Stimulation of dSPN_ALM_ in the right hemisphere during the sample or delay epoch biased upcoming licking from the right to left spouts (Figure S3B), an opposite directional bias to stimulation in the left hemisphere. Stimulation of iSPN_ALM_ caused an opposite directional bias to stimulation of dSPN_ALM_. Thus, stimulation of the direct (indirect) pathway in the left and right striatum caused opposite behavioral bias, despite that dSPN_ALM_ and iSPN_ALM_ in both hemispheres were anterogradely labelled from the same cortical region.

### Similar response pattern in dSPN_ALM_ and iSPN_ALM_ during behavior

Striatal dSPN_ALM_ and iSPN_ALM_ played opposing roles in STM (Figure 1), suggesting that the two types of neurons respond differently in the decision task. To check this, we need to perform cell-type specific neurophysiology. As dSPNs and iSPNs have similar firing rates and spiking waveforms, they cannot be reliably separated based on these neurophysiological properties. Thus, we performed optogenetic tagging to identify ALM-input defined dSPNs and iSPNs (Shin et al., 2018). To increase the efficiency of tagging, we customized an optrode with two optical fibers attached above recording sites (Figure 2). We then advanced the optrode to the ALM projection zone of the striatum in transgenic mice (R26 LSL FSF ReaChR-mCitrine x D1-cre or D2-cre) with anterograde transsynaptic virus (AAV1-hSyn-FLPO) injected in ALM (Figure 2A). Laser power was properly tuned as we found that high laser power could lead to a detrimental effect in tagging efficiency, probably caused by stronger local inhibition (Figure 2F). We identified 1769 single-units that were reliably activated by laser stimulation with short latency (significant increase within 6 ms, P < 0.01, t-test, 624 single-units from 7 D1-Cre and 1145 single-units from 6 D2-Cre mice, Figure 2B, C). Among these single-units, we selected 467 putative dSPN_ALM_ and 914 putative iSPN_ALM_ that had high waveform similarity (Figure 2D, E, criterion see Methods). Selected dSPN_ALM_ and iSPN_ALM_ had a similar latency upon laser stimulation (4.64 ± 0.74 ms vs 4.71 ± 0.72 ms, Mean ± SD). A considerable fraction of tagged neurons exhibited very low firing rate (< 0.5 Hz, corresponding to 1 spike during the STM epoch in every two trials, threshold arbitrarily chosen, Figure 2H, I). There were more quiet dSPN_ALM_ than iSPN_ALM_ (47.4 ± 13.0% vs 26.3 ± 3.2%, P = 0.0027, t-test, Figure S4G). We hereafter focused on non-quiet neurons if not explicitly specified.

**Fig. 2.**
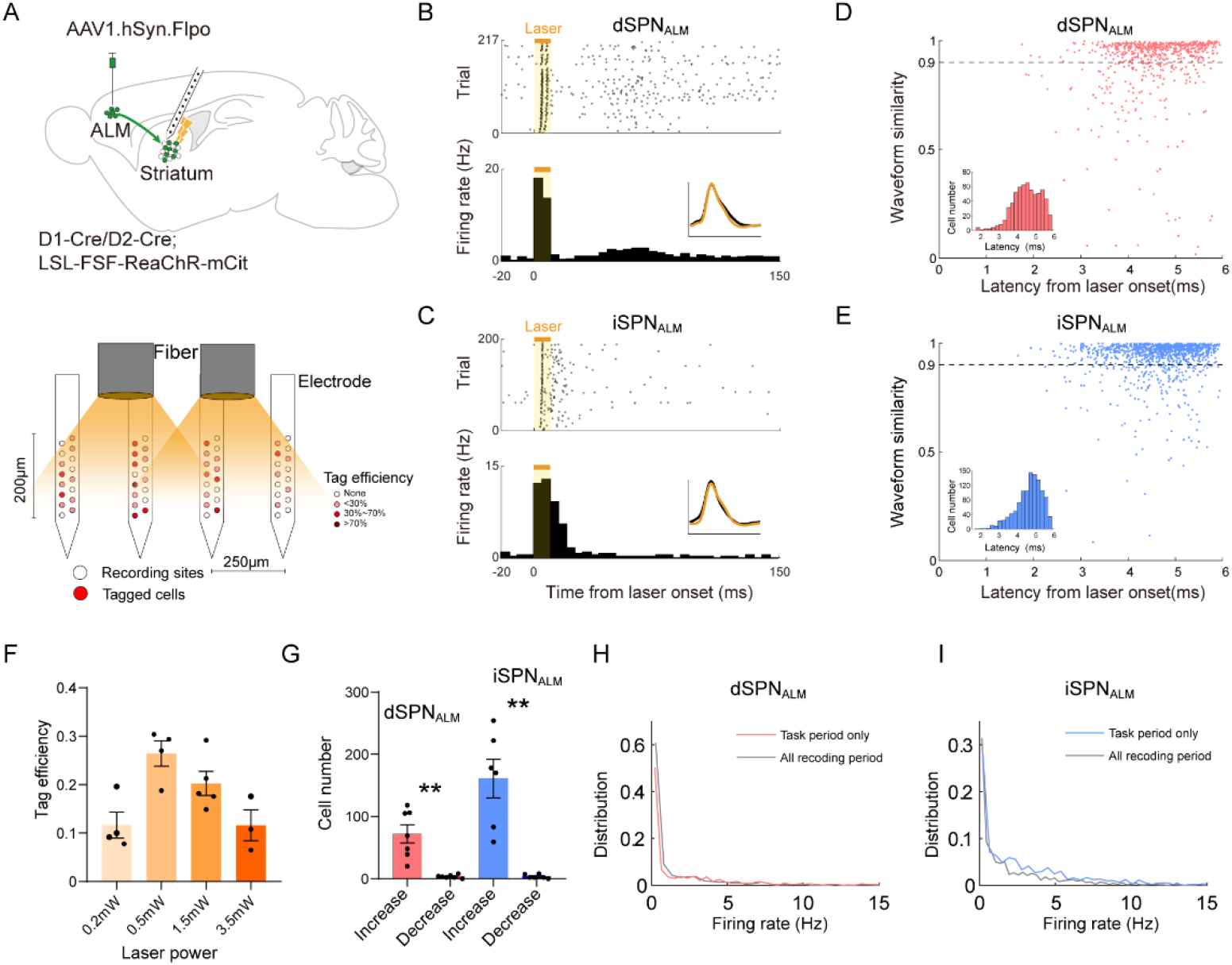
Cell-type specific recording of dSPN_ALM_ and iSPN_ALM_. (A) Top, schematic of identification of dSPN_ALM_ or iSPN_ALM_. After each behavioral session, a train of pulses were delivered to activate putative dSPN_ALM_ or iSPN_ALM_ that expressed ReaChR. Bottom, schematic of a 64-channel silicon probe with two 100-μm-diameter fibers attached above recording sites. Colored dots indicate tag efficiency of recording sites from 11 behavioral sessions. (B) A putative dSPN_ALM_ neuron. Top: spike raster. Each dot represents one spike and each row represents one stimulation trial. The orange bar indicates the stimulation period (10 ms). Bottom, peri-stimulus time histogram (PSTH). Binsize, 5 ms. Inset, average waveforms of spontaneous (black) and light-evoked (orange) spikes. (C) A putative iSPN_ALM_ neuron. (D) Waveform similarity between spontaneous and light-evoked spikes of all tagged dSPN_ALM_ neurons. Inset, distribution of spike latency relative to laser onset (Mean ± SD: 4.64 ± 0.74 ms). (E) Same format as in (D) but for tagged iSPN_ALM_ neurons. Spike latency, 4.71 ± 0.72 ms (Mean ± SD). (F) Fraction of tagged cells under different laser powers. Each dot represents one recording session in one representative mouse with iSPN_ALM_ expressing ReaChR. (G) The number of significantly modulated neurons. Each dot represents one mouse (n = 7 D1-Cre mice, n = 6 D2-Cre mice). (H) Firing rate distribution of dSPN_ALM_ within the task period (light red) and during the whole recording period (gray). (I) Firing rate distribution of iSPN_ALM_. ** denotes log-rank test with statistical significance P < 0.01.

Activity of a large fraction of neurons in both direct and indirect pathways differentiated trial types (33.4%, 156/467 in dSPN_ALM_; 45.8%, 419/914 in iSPN_ALM_; Methods, see example neurons in Figure 3A, B, and summary in Figure 3C, D). Neural responses in both pathways were diverse: subsets of neurons showed selective sample and/or delay activity (13.3%, 62/467 in dSPN_ALM_; 20.9%, 191/914 in iSPN_ALM_), peri-movement activity during the response epoch (12.4%, 58/467 in dSPN_ALM_; 11.7%, 107/914 in iSPN_ALM_), or both (4.1%, 19/467 in dSPN_ALM_; 4.0%, 37/914 in iSPN_ALM_, Figure 3C, D). In both pathways, there were approximately equal numbers of neurons preferred contra- or ipsi-trials during delay or response epochs (not significant, t-test, Figure S4E, F). More SPNs preferred contra-rather than ipsi-trials in the sample epoch in either pathway (11.8 ± 4.8% vs 1.5 ± 2.1% in dSPN_ALM_, P < 0.001, t-test; 20.8 ± 8.8% vs 3.4 ± 3.0% in iSPN_ALM_, P = 0.001, t-test, Figure S4E, F). The higher contra-preference in the sample epoch was probably caused by the stronger tactile stimulus as many of these neurons responded similarly in error trials (i.e. sensory-related, Figure 3A, B, G, H). During the delay and response periods, many neurons switched their responses in correct and error trials, indicating that activity in these neurons closely tracked animals’ choice (Figure 3A, B, G, H). The mean activity of dSPN_ALM_ and iSPN_ALM_ evolved in time with similar patterns for non-quiet neurons (4.81 ± 0.41 vs 4.57 ± 0.18 Hz during the task period, Mean ± SEM, P = 0.5498, t-test, Figure 3E), although there existed significant difference in firing rates if taking all tagged SPNs into consideration (2.62 ± 0.25 vs 3.44 ± 0.15 Hz during the task period, Mean ± SEM, P < 0.01, t-test, Figure 2H, I). Selectivity (defined as the firing rate difference in contra- and ipsi-trials) also showed a similar pattern with time during the delay and response epoch, although the magnitude was higher in the indirect pathway (Figure 3F). Thus, SPNs of the direct and indirect pathways show similar response patterns during the tactile-based decision making task.

**Fig. 3.**
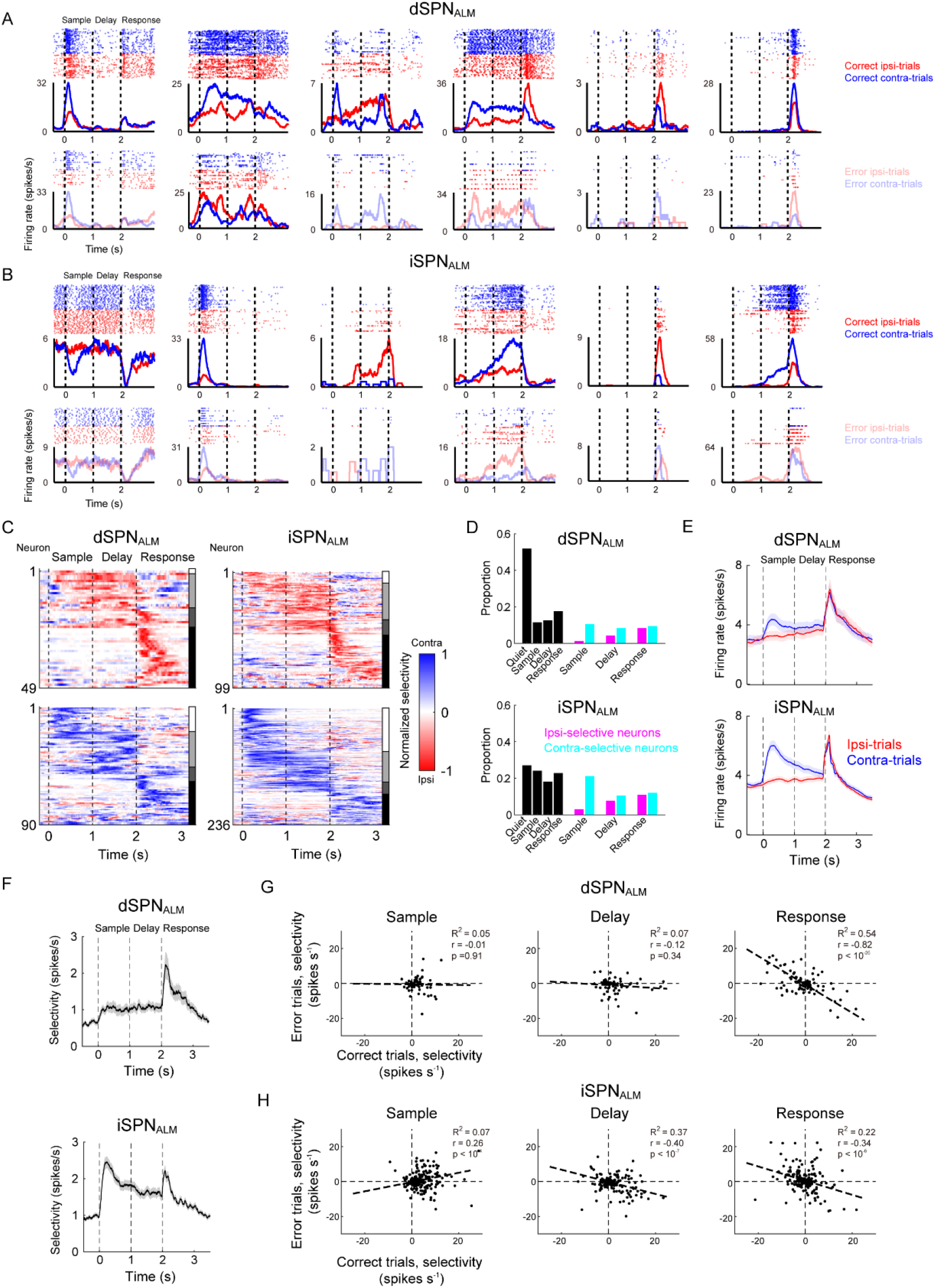
Activity of dSPN_ALM_ and iSPN_ALM_ during the decision task. (A) Activity of six example dSPN_ALM_ neurons during correct (top) and error trials (bottom). Red, ipsi-trials; Blue, contra-trials. Dashed lines separate behavioral epochs. Bin size, 1 ms. Smooth window, 100 ms. (B) Similar to (A) but for iSPN_ALM_. (C) Population selectivity of dSPN_ALM_ (left) and iSPN_ALM_ (right). Red, ipsilateral selective neurons. Blue, contralateral selective neurons. Vertical bars on the right: white, neurons that are selective only in the sample epoch; light grey, neurons with selectivity in the delay and/or sample epoch; dark grey, neurons with both sample/delay and peri-movement activity; black, neurons with peri-movement activity only. Units switched preference in different epochs were excluded. (D) Proportion of quiet and selective dSPN_ALM_ (top) and iSPN_ALM_ (bottom) during the sample, delay and response epochs. Magenta, ipsi-selective. Cyan, contra-selective. Black, ipsi- and contra-selective neurons combined. (E) Mean activity of dSPN_ALM_ (top) and iSPN_ALM_ (bottom) during the tactile-based decision task. Red, ipsi-trials. Blue, contra-trials. Shadow, SEM. Smooth window, 200 ms. Quiet cells were excluded. (F) Mean absolute selectivity of dSPN_ALM_ (left) and iSPN_ALM_ (right). Here the absolute selectivity was calculated as the spike rate in preferred trials subtracted with that in non-preferred trials. Shadow, SEM. Smooth window, 200 ms. (G) Selectivity of dSPN_ALM_ in correct and error trials during sample (left), delay (middle) and response epochs (right). Each dot represents a single neuron. Selectivity was calculated as the spike rate difference between the contra- and ipsi-trials. Only selective neurons during each period were included. Dotted line, linear regression. (H) Same format as in (G) but for iSPN_ALM_.

As ALM-input defined SPNs spanned a large region in striatum, we examined whether SPNs with different functional responses had distinct spatial organization. To localize recording sites, we painted the silicon probe with a thin layer of DiI and applied a brief electric pulse to mark a small lesion near the tip of the probe (Huo et al., 2020). We then imaged the whole mouse brain using a multiscale light-sheet microscope (Zhang et al., 2021) and mapped the electrode tracks as well as the lesion location to the common coordinate framework (Figure 4A). This pipleline has been verified with electrophysiology landmarks and has an accuracy up to ~20 μm (Zhang et al., 2021). Optogenetically identified dSPN_ALM_ and iSPN_ALM_ were distributed widely in the central-ventral striatum, and different response-types of neurons (contra- or ipsi-preferring neurons) were spatially mixed (Figure 4E, F). Striatal dSPN_ALM_ and iSPN_ALM_ within ~100 μm showed a high correlation in activity (Figure 4C), consistent with the finding that striatal neurons are organized in functional clusters to represent movement and reward information (Barbera et al., 2016; Shin et al., 2020).The correlation decayed faster in dSPN_ALM_ compared with iSPN_ALM_ (Figure 4C). This was not caused by the larger proportion of quiet dSPN_ALM_ as the result was similar for non-quiet neurons (Figure S5C). We further examined how selectivity in dSPN_ALM_ and iSPN_ALM_ evolved over time. To do so, we selected eight epochs (each lasting 200ms; the baseline, early sample, late sample, early delay, late delay, early response, late response and trial interval epochs) and projected selectivity of recorded neurons along four coronal slices (Figure 4E, F). There were more contra-preferring neurons in the sample epoch in both pathways, consistent with previous analysis (Figure 3D, S4E, F). More ipsi-preferring neurons emerged starting from the late delay epoch and reached a peak during the response epoch (Figure 4E, F). This pattern was consistently observed across multiple projection planes and it was more evident in iSPN_ALM_. Interestingly, the fraction of ipsi-preferring neurons reached its peak in the early response epoch for iSPN_ALM_ while it only summited in late response epoch for dSPN_ALM_ (Figure 4D-F; Figure S5A). This pattern cannot be explained by difference in licking response as there was no difference in first-lick latency (Figure S5B). Thus, despite that dSPN_ALM_ and iSPN_ALM_ showed overall similar response patterns, they had distinct spatiotemporal organization in the decision task.

**Fig. 4.**
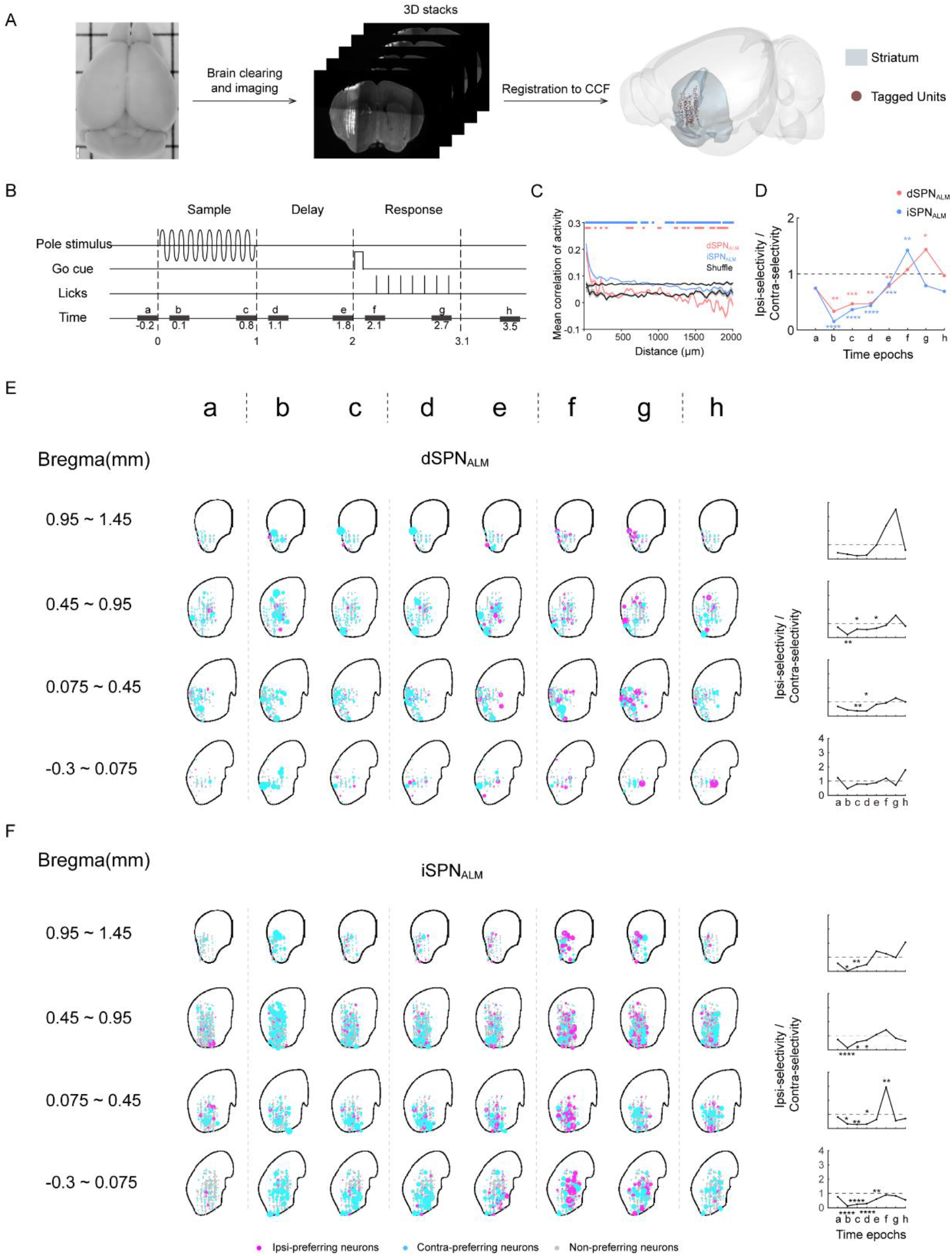
Spatiotemporal distribution of dSPN_ALM_ and iSPN_ALM_ activity. (A) Pipeline of 3D reconstruction of recording sites. (B) Time windows (0.2s) chosen for temporal analysis of neural activity. (C) Correlation of activity in both dSPN_ALM_ (light red) and iSPN_ALM_ (light blue) decays with increased distance between pairs of neurons. However, correlation in dSPN_ALM_ decays at a faster rate than that in iSPN_ALM_. Black, Shuffled data. Window size, 50ms. All neurons included. Significance from shuffled data was indicated by colored bars above (two-sided t-test, P < 0.01). (D) Summed selectivity ratio (see Methods) of dSPN_ALM_ and iSPN_ALM_ in eight time epochs. Ipsi-preference increases earlier in iSPN_ALM_ compared with dSPN_ALM_. (E) Spatiotemporal distribution of selectivity in dSPN_ALM_ (magenta, ipsi-preferring neurons; cyan, contra-preferring neurons). More ipsi-preferring neurons emerged in the response epoch. The size of dots symbolizes the magnitude of selectivity. Each row shows neurons projected on the indicated coronal slice. Right panel, summed selectivity ratio during different time epochs for neurons on the corresponding slice. (F) Same format as in (E) but for iSPN_ALM_ neurons. *, **, *** & **** denote two-sided t-test with statistical significance P < 0.05, P < 0.01, P < 0.001& P < 0.0001, respectively.

### Direct and indirect pathway SPNs specifically modulate ALM activity

Stimulation of dSPN_ALM_ and iSPN_ALM_ oppositely biased upcoming choice while the two types of neurons have similar response patterns during the tactile-based decision task. As population activity in ALM predicts upcoming choice on a trial-by-trial basis (Li et al., 2016), we studied how stimulation of dSPN_ALM_ and iSPN_ALM_ regulates ALM population activity (Figure 5). As stimulation of dSPN_ALM_ produced a contralateral bias that was similar to activation of ALM pyramidal tract neurons directly, we hypothesized that stimulation of dSPN_ALM_ would increase ALM activity. To test this, we recorded ALM population activity with simultaneous perturbation of the direct pathway during the decision task (Figure 5A). We first focused on putative pyramidal neurons (pPNs, n = 756/908 from 7 mice) (see putative fast spiking neurons [pFSNs] in Figure S6, Methods). Contrary to our simple hypothesis, stimulation of dSPN_ALM_ during the delay epoch did not uniformly increase ALM activity (see example neurons in Figure 5B). There were similar fractions of up- and down-modulated neurons (mean firing rate significantly changed, 14.3% up, 12.6% down, units selected using t-test with P < 0.05, Figure 5E; see Figure S7 for activity during sample epoch stimulation). Interestingly, neurons with different response properties were modulated differentially. The delta activity, defined as the activity difference between stimulation and control trials, was significantly larger in contra-preferring neurons than in ipsi-preferring neurons (P < 0.05, t-test, Figure 5E). The differential effect was caused by the larger impact on ipsi-rather than conta-trials (Figure 5G, left; example neurons in Figure 5B), with activity in contra-preferring neurons was increased on ipsi-trials and activity in ipsi-preferring neurons was decreased on ipsi-trials (Figure 5H). These results indicate that stimulation of dSPN_ALM_ regulates ALM activity differently in different response-types of neurons.

**Fig. 5.**
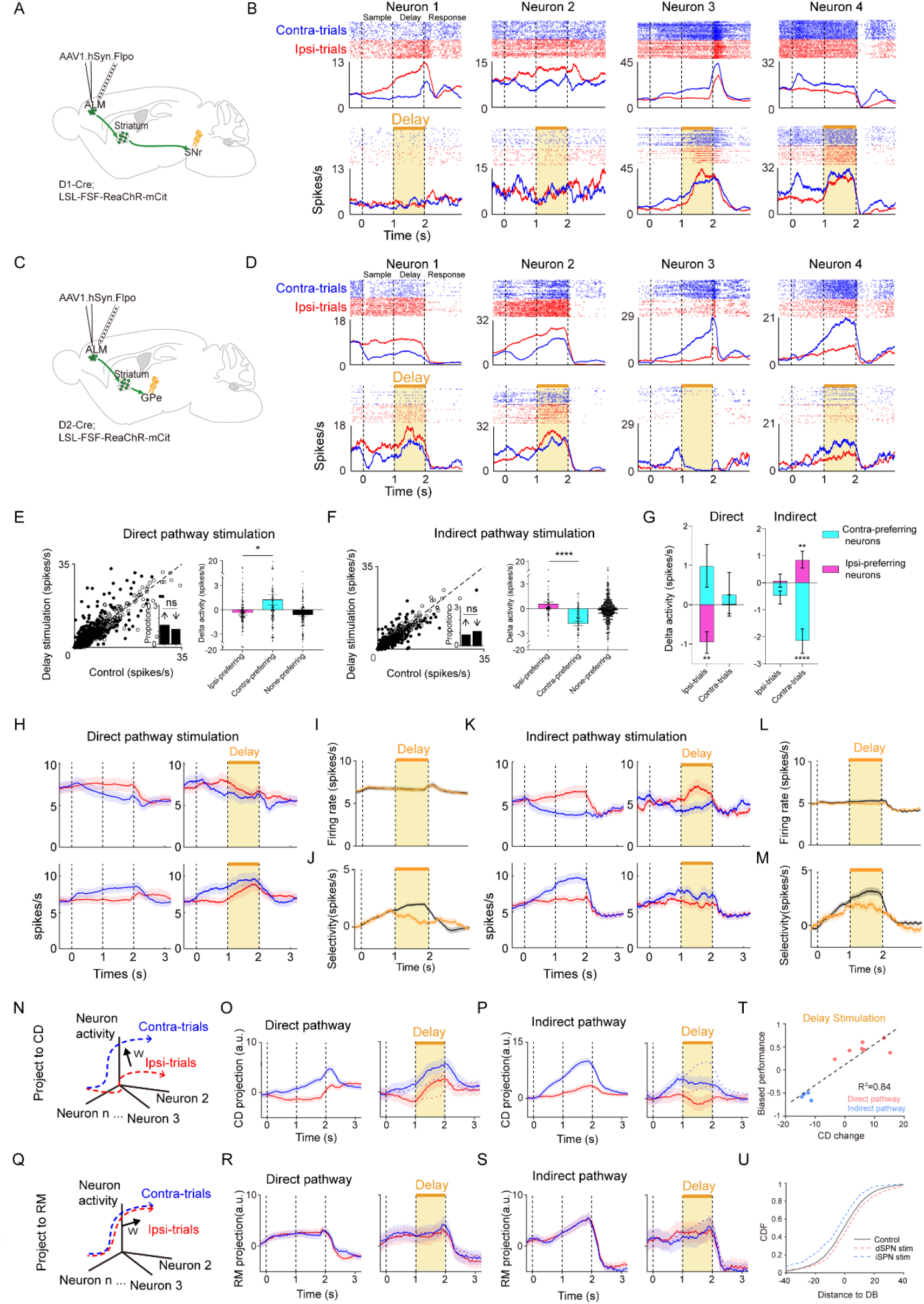
Stimulation of ALM-input defined direct and indirect pathways oppositely modulate ALM activity. (A) Schematic of ALM recording with simultaneous stimulation of dSPN_ALM_ by illuminating their axons in SNr. (B) Four example ALM neurons during stimulation of the direct pathway. Top: spike raster and PSTH. Bottom: spike raster and PSTH during stimulation of the direct pathway. Dashed lines separate behavioral epochs. Bin size, 1ms. Averaging window, 100ms. (C) Schematic of ALM recording with simultaneous stimulation of iSPN_ALM_ by illuminating their axons in GPe. (D) Four example ALM neurons during stimulation of the indirect pathway. Same format as in (B). (E) Scatter plot of mean firing rates during the delay epoch (756 units from 7 mice). Filled circles, neurons that were significantly modulated (P < 0.05, t-test). Inset, fraction of up- and down-modulated neurons (no significant difference, Chi-square test). Dotted line is the unity line. Right, delta activity for ipsi-preferring, contra-preferring and non-preferring neurons. (F) Same format as in (E) but for stimulation of the indirect pathway (624 units from 4 mice). (G) Comparison of delta activity between ipsi- and contra-trials. Stimulation of the direct pathway (left) caused large activity change on ipsi-trials for both contra-preferring and ipsi-preferring neurons, while stimulation of the indirect pathway (right) caused large activity change on contra-trials. (H) Activity of selective neurons during direct pathway stimulation. Top, ipsi-selective neurons (n= 129). Bottom, contra-selective neurons (n = 92). Left, control trials. Right, delay stimulation. (I) Mean PSTH of ALM neurons during control (black) and stimulation (orange) of the direct pathway (n = 756 neurons). Shading, SEM. (J) Selectivity of ALM neurons during control (black) and stimulation (orange) of the direct pathway. (K) Same format as in (H) but for indirect pathway stimulation. Ipsi-selective neurons, n = 67. Contra-selective neurons, n = 92. (L) Mean PSTH of ALM neurons during control (black) and stimulation (orange) of the indirect pathway (N = 624 neurons). Shading, SEM. (M) Selectivity of ALM neurons during control (black) and stimulation (orange) of the indirect pathway. (N) Schematic of population activity projected along the coding direction (CD). Blue and red curves indicate the population trajectory on contra- and ipsi-trials, respectively. W, coding direction. (O) CD-projected activity during control (left) and stimulation (right) of the direct pathway. Shading, SEM (n = 7 mice). Bin size, 20 ms. Averaging window, 400 ms. (P) Same format as in (O) but for stimulation of indirect pathway. (Q) Schematic of population activity projected along the ramping mode (RM). (R) RM-projected activity during control (left) and stimulation (right) of the direct pathway. Same data as in (O). (S) Same format as in (R) but for stimulation of indirect pathway. (T) Linear correlation between behavioral bias and CD change. Each circle represents a mouse. Red, direct pathway stimulation; Blue, indirect pathway stimulation. (U) Cumulative distribution function (CDF) of distance to the decision boundary (DB) during control (black), direct pathway (red) and indirect pathway (blue) stimulation conditions. *, **, & **** denote two-sided t-test with statistical significance P < 0.05, P < 0.01, & P < 0.0001, respectively. ns, not significant.

We further checked whether stimulation of the indirect pathway specifically affected ALM activity. As stimulation of iSPN_ALM_ produced an ipsilateral bias that was similar to inactivation of ALM, we hypothesized that stimulation of iSPN_ALM_ would decrease ALM activity. Contrary to the hypothesis, stimulation during the delay epoch did not uniformly decrease ALM activity either (see example neurons in Figure 5D). There were similar fractions of up- and down-modulated neurons (mean firing rate significantly changed, 9.1% up, 11.5% down, units selected using t-test with P < 0.05, Figure 5F; see Figure S7 for activity during sample epoch stimulation). Close examination indicated that neurons with different response properties were differentially modulated. The delta activity was significantly lower in contra-preferring neurons than in ipsi-preferring neurons (P < 0.0001, t-test, Figure 5F). The differential effect was caused by the larger impact on contra-rather than ipsi-trials (Figure 5G, right; example neurons in Figure 5D and averaged activity in Figure 5K). Thus, stimulation of iSPN_ALM_ also specifically regulates ALM activity, but different from stimulation of dSPN_ALM_, mainly on contra-trials.

Overall, stimulation of dSPN_ALM_ or iSPN_ALM_ affected ALM mean activity little but significantly reduced selectivity (defined as the absolute difference of activity in contra- and ipsi-trials; selectivity reduction, 64.2 ± 12.5% during the last 300 ms of the delay epoch for dSPN_ALM_, Mean ± SEM, t-test, P < 0.001; 41.7 ± 10.4% for iSPN_ALM_, P = 0.0049; Figure 5I, J, L, M). Contrary to pPNs, pFSNs were more uniformly affected by stimulation of the direct and indirect pathways (stimulation of dSPN_ALM_, increased vs decreased, 21% vs 8.4%; stimulation of iSPN_ALM_, 6.8% vs 17.8%, Figure S6). These results indicate that the direct and indirect pathways modulate ALM pyramidal neurons in a response-type and trial-type specific way while the effect on fast spiking neurons was less specific.

To understand how stimulation of dSPN_ALM_ and iSPN_ALM_ modulated ALM population activity to produce opposite behavioral deficits, we examined simultaneously recorded neurons in the activity space. We focused on pPNs because there were few simultaneously recorded pFSNs. Population activity in ALM on contra- and ipsi-trials evolved with trial progression to form distinct trajectories (Li et al., 2016). We then performed dimensionality reduction by projecting trajectories along the coding direction (CD, along which activity maximally discriminated upcoming choices), and along the Ramping Mode (RM), which captured the non-specific increase of activity (Figure 5N, Q). Stimulation of dSPN_ALM_ terminals shifted the CD-projected trajectories in the ipsi- to contra-direction (Figure 5O), while stimulation of iSPN_ALM_ shifted the trajectories in the opposite direction (Figure 5P), consistent with the opposite behavioral deficits (Figure 1). Stimulation of both pathways affected projections along the ramping mode little (Figure 5R, S). As CD-projected activity predicts movement direction on a trial-by-trial basis (Li et al., 2016), activity along CD represents a decision variable according to which mice decide which lick-spout to choose. Indeed, the magnitude of shift along CD caused by stimulation was linearly correlated with the degree of behavioral bias (Figure 5T). And stimulation of direct and indirect pathways bi-directionally shifted the distribution of CD-projected activity (Figure 5U, see Method). These results indicate that the direct and indirect pathways function in tug-of-war to properly shape cortical trajectories underlying STM.

### Direct and indirect pathway SPNs oppositely regulate SNr ramping activity

Striatal dSPN_ALM_ and iSPN_ALM_ oppositely regulate ALM trajectories, likely through their effect on the basal ganglia output nucleus SNr that projects to ALM-reciprocally connected thalamic nuclei (Guo et al., 2017). To check where the opposing effects first appear along the basal ganglia-thalamocortical pathway, we recorded SNr population activity during stimulation of dSPN_ALM_ and iSPN_ALM_. As dSPN_ALM_ neurons are GABAergic, we hypothesized that stimulation of dSPN_ALM_ would uniformly reduce SNr activity. To test this, we implanted an optrode in SNr to both activate dSPN_ALM_ terminals and record SNr activity (Figure 6A). We selected putative GABAergic neurons (n = 215/303 from 6 mice) based on their narrow spike waveforms (criterion previously verified by optical tagging (Wang et al., 2021), Methods). Contrary to our hypothesis, stimulation of dSPN_ALM_ during the delay epoch did not uniformly reduce SNr activity (see example neurons in Figure 6B). Although a large fraction of neurons displayed decreased activity (30.9%, neurons selected using t-test using significance P < 0.05), a significant fraction of neurons increased their activity (16.2%) (Figure 6C). Striatal iSPNs can increase SNr activity indirectly through the external globus pallidus (GPe) and the subthalamic nucleus (STN). However, stimulation of iSPN_ALM_ did not uniformly increase SNr activity either (Figure 6D, E). A large fraction of neurons displayed modulated activity (mean firing rate significantly changed, 17.7% up, 11.8% down, Figure 6F).

**Fig. 6.**
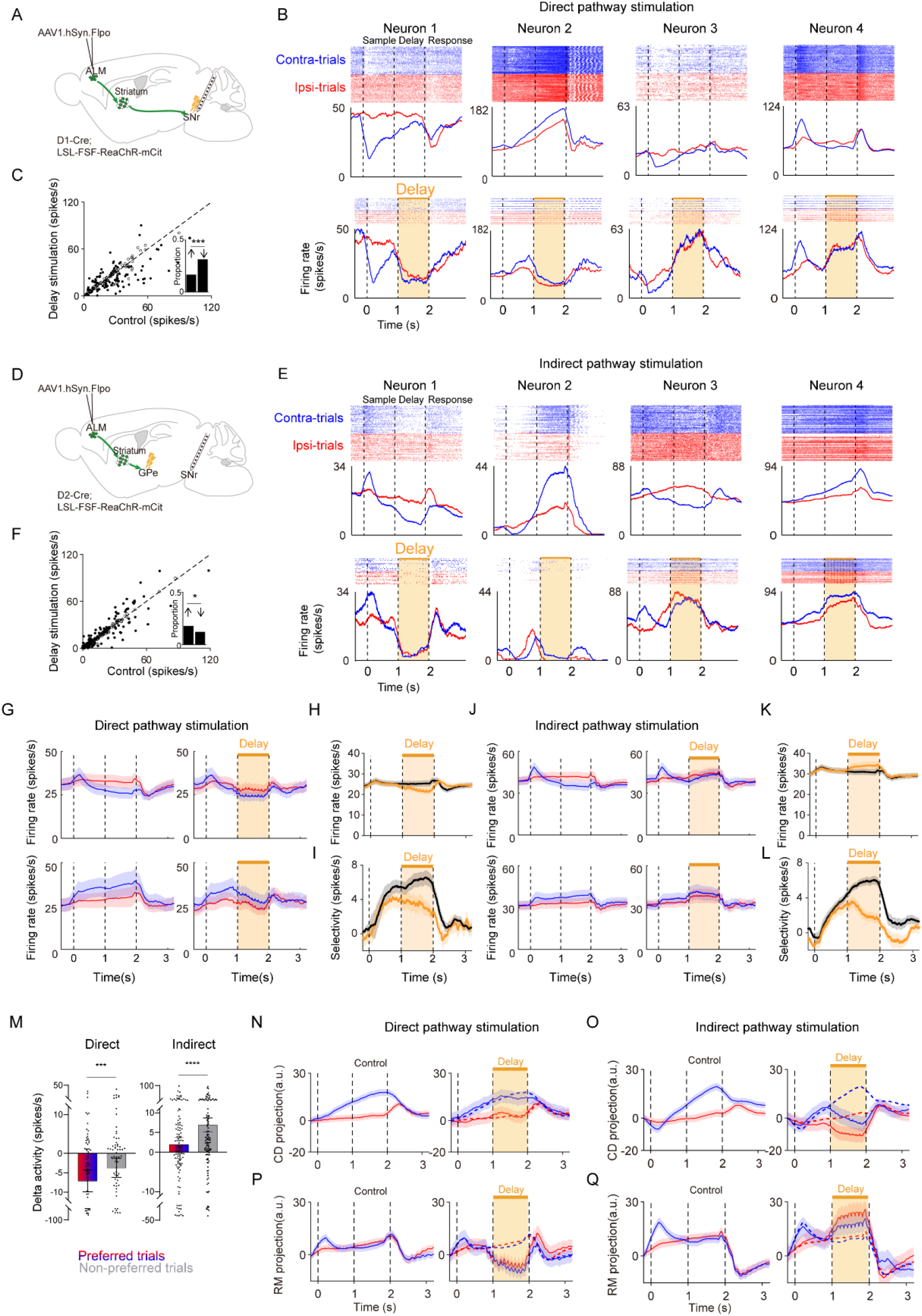
Stimulation of ALM-input defined direct and indirect pathways differentially modulate SNr activity. (A) Schematic of SNr recording with simultaneous stimulation of axons of dSPN_ALM_. (B) Four example SNr neurons during stimulation of the direct pathway. Top: spike raster and PSTH. Bottom: spike raster and PSTH during stimulation of the direct pathway. Dashed lines separate behavioral epochs. Bin size, 1ms. Averaging window, 100ms. (C) Scatter plot of mean firing rates during the delay epoch (215 units from 6 mice). Filled circles, neurons that were significantly modulated (P < 0.05, t-test). Dotted line is the unity line. Inset, proportion of up-modulated and down-modulated neurons. (D) Schematic of SNr recording with simultaneous stimulation of axons of iSPN_ALM_. (E) Four example SNr neurons during stimulation of the indirect pathway. (F) Same format as in (C), but for SNr neurons during stimulation of the indirect pathway (n = 323). (G) Mean PSTH of SNr neurons during control (left) and stimulation (right) of dSPN_ALM_. Shading, SEM. Top, ipsi-preferring neurons (n = 40) selected based on delay epoch activity; bottom, contra-preferring neurons (n = 24 neurons). Red, ipsilateral trials; blue, contralateral trials. (H) Mean PSTH of SNr neurons during control (black) and stimulation (orange) of the direct pathway (n = 215 neurons). Shading, SEM. (I) Selectivity of SNr neurons during control (black) and stimulation (orange) of the direct pathway. (J) Same format as in (G) but for stimulation of the indirect pathway (ipsi-preferring neurons, n = 53; contra-preferring neurons, n = 58). (K) Mean PSTH of SNr neurons during control (black) and stimulation (orange) of the indirect pathway (n = 323 neurons). Shading, SEM. (L) Selectivity of SNr neurons during control (black) and stimulation (orange) of the indirect pathway. (M) Activity change in preferred trials (red-blue gradient color) or non-preferred trials (gray) during stimulation of dSPN_ALM_ (left) or iSPN_ALM_ (right). Delay-selective neurons were analyzed (dSPN_ALM_: n = 64; iSPN_ALM_: n = 111). (N) CD-projected activity during control (left) and stimulation of the direct (right) pathways. Shading, SEM (n = 6 mice). Bin size, 20 ms. Averaging window, 20 ms. (O) Same format as in (N) but for stimulation of the indirect pathway (n = 7 mice). (P) RM-projected activity during control (left) and stimulation of the direct (right) pathways. Same data as in (N). (Q) Same format as in (P) but for stimulation of the indirect pathway. Same data as in (O). * & *** in C, F denote Chi-square test with statistical significance P < 0.05 & P < 0.001, respectively. *** & **** in M denote paired two-sided t-test with statistical significance P < 0.001 & P < 0.0001, respectively.

Overall, stimulation of dSPN_ALM_ reduced SNr mean activity by 14.4 ± 3.8% (calculated during the last 300 ms of the delay epoch, Mean ± SEM, paired t-test, P < 0.001, Figure 6H) but dramatically reduced selectivity by 48.5 ± 13.2% (paired t-test, P < 0.001, Figure 6I). Stimulation of iSPN_ALM_ increased SNr mean activity by 10.2 ± 2.5% (paired t-test, P < 0.0001, Figure 6K) and dramatically reduced selectivity by 74.2 ± 10.4% (paired t-test, P < 0.0001, Figure 6L). Close examination indicated that stimulation of dSPN_ALM_ and iSPN_ALM_ affected SNr activity differently in different trial-types (Figure 7). Stimulation of dSPN_ALM_ decreased SNr activity mainly in preferred trials (contra-trials in contra-preferring neurons and ipsi-trials in ipsi-preferring neurons, population activity in Figure 6G, individual neurons in Figure 6M, delta activity in preferred trials vs non-preferred trials, −7.04 ± 2.83 vs −3.69 ± 2.45 Hz, Mean ± SEM, paired t-test, P < 0.001). On the contrary, stimulation of iSPN_ALM_ increased SNr activity mainly in non-preferred trials (delta activity in preferred trials vs non-preferred trials, 1.98 ± 1.71 vs 6.94 ± 1.76 Hz, Mean ± SEM, paired t-test, P < 0.0001, Figure 6J, M). These results indicate that stimulation of the direct and indirect pathways affect SNr activity in a trial-type dependent way.

**Fig. 7.**
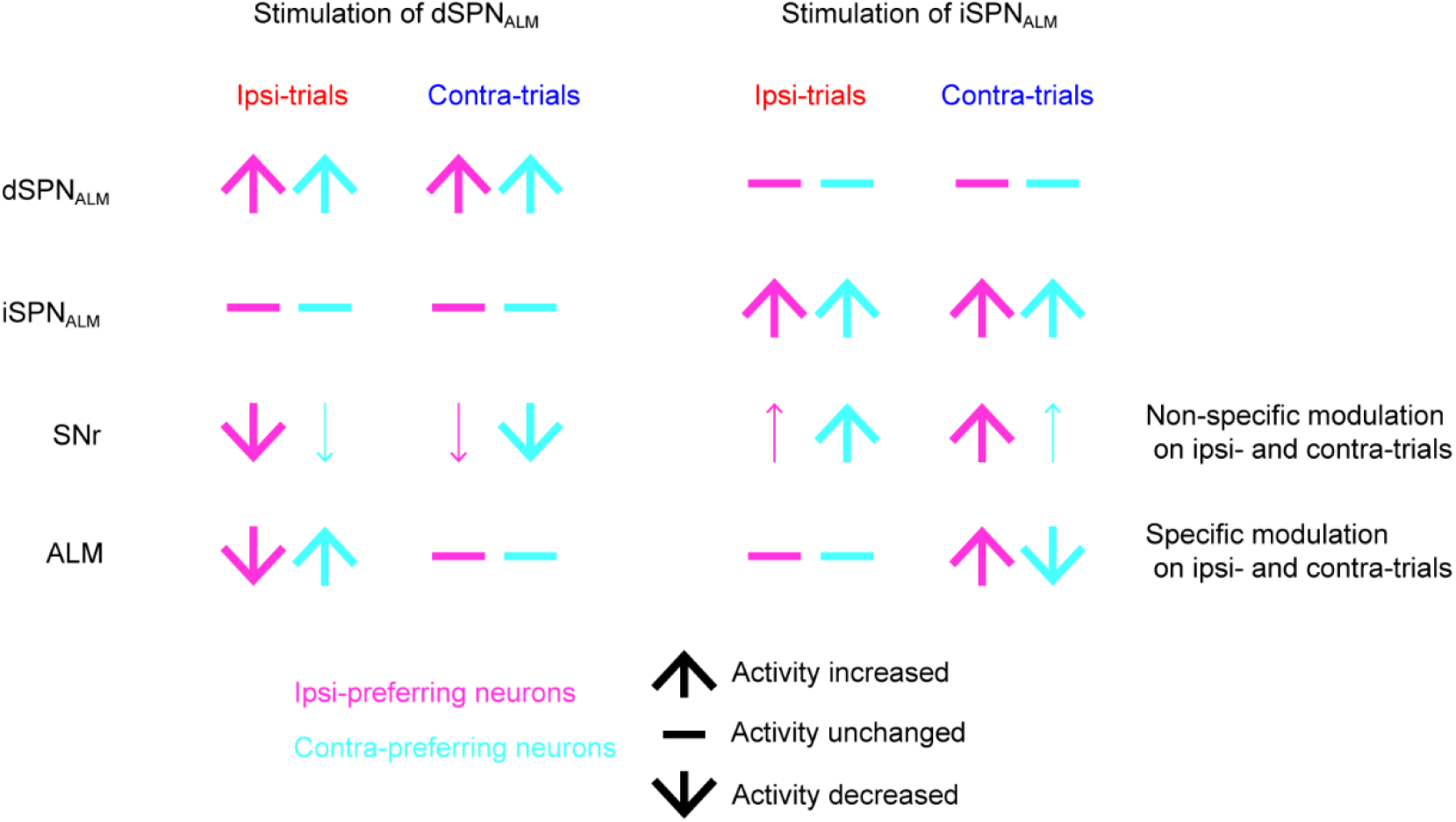
Summary of effects of stimulation of direct and indirect pathways on activity in SNr and ALM. The modulation of SNr depends on trial-types. The modulation of ALM depends on trial-types and response-types.

To understand the diverse effect of perturbation on individual SNr neurons, we projected neural trajectories along the CD and RM directions (Figure 6N-Q). Activity along CD did not explain the opposite pattern of behavioral deficits as the effect due to stimulation of dSPN_ALM_ was minimal and in the same direction along that caused by stimulation of iSPN_ALM_ (Figure 6N, O). Along RM, stimulation of dSPN_ALM_ and iSPN_ALM_ reduced and increased SNr activity, respectively (Figure 6P, Q). The modulation direction along RM was consistent with the anatomical projections of the direct and indirect pathway SPNs. The opposite effects of stimulation on RM-projected trajectories suggest that non-specific perturbation of SNr activity leads to opposite behavioral deficits observed in the decision task. Consistently, non-selective activation and inactivation of SNr-to-thalamus projections caused an ipsilateral and contralateral bias, respectively (Wang et al. 2021). These results indicate that the direct and indirect pathways oppositely modulate cortical activity by differentially regulating SNr ramping activity.

## Discussion

Many studies found that the direct and indirect pathways played antagonistic roles in regulating movement while direct and indirect pathway SPNs had concurrent activation during movement (Barbera et al., 2016; Chen et al., 2021; Cui et al., 2013; Jin et al., 2014; Kravitz et al., 2010; Lee and Sabatini, 2021; Markowitz et al., 2018; Oldenburg and Sabatini, 2015; Tecuapetla et al., 2016). We verified previous findings by recording and perturbing dSPNs and iSPNs in the same well-controlled decision task (Figure 1, 3). Previous studies typically focus on regions of the striatum based on spatial divisions and study movements that involves complicated kinetics. We here specifically labelled dSPNs and iSPNs that received synaptic input from ALM, whose role in STM has been well established, and studied their role in a quasi-steady state (delay period) (Figure 1). Even with this specificity, we still found that stimulation of the direct and indirect pathways oppositely biased upcoming choice (Figure 1). And cell-type specific recordings showed that direct and indirect pathway SPNs had overall similar response patterns during the STM period (Figure 3). These results suggest that the direct and indirect pathways also have opposing roles in cognition, and the opposing roles are due to their concurrent activation.

Genetic and optogenetic perturbation of dSPNs and iSPNs oppositely modulate locomotor activity, lever-pressing, licking as well as reinforcement learning (Chen et al., 2021; Kravitz et al., 2010; Lee and Sabatini, 2021; Oldenburg and Sabatini, 2015; Tai et al., 2012; Tecuapetla et al., 2016; Yttri and Dudman, 2016). We demonstrated that stimulation of the direct and indirect pathways during the STM period oppositely biased upcoming choice, extending the classic model to the domain of cognition. We note that, although our results are consistent with the classic model, stimulation of dSPN_ALM_ and iSPN_ALM_ using high laser powers affected licking differently during the supposed no-licking period (Figure S2A, B), with stimulation of iSPN_ALM_ inducing early-licks while stimulation of dSPN_ALM_ having no effect. Consistently, strong stimulation of direct pathway neurons suppressed, rather than facilitated, lever-pressing (Tecuapetla et al. 2016). These results are somewhat difficult to reconcile if the direct pathway simply has a prokinetic role while the indirect pathway has an antikinetic role. Furthermore, it is noted that stimulation of either pathway before sequence initiation increases the latency for level-pressing initiation (Tecuapetla et al. 2016), and promotes “yes” responses in a visual detection task (Wang et al. 2018). As the striatum is composed of multiple domains with diverse cortical inputs (Hintiryan et al., 2016; Hunnicutt et al., 2016) and striatal neurons seem to be composed of functionally distinct spatial clusters (Barbera et al., 2016; Shin et al., 2020), modulating functional-specific dSPNs and iSPNs clusters with defined cortical input can potentially reveal the precise mechanisms by which the direct and indirect pathways regulate movement.

The direct and indirect pathway SPNs have been found to be co-active during movement initiation and termination (Barbera et al., 2016; Cui et al., 2013; Jin et al., 2014; Markowitz et al., 2018). With optogenetic identification of ALM-input defined dSPNs and iSPNs, we further found that the direct and indirect pathways were concurrently active during STM (Figure 3). It is noted that, although dSPNs and iSPNs have coarse pattern of co-activation, there exist quantitative difference in firing patterns to support behavioral variables related to movement, reward value, reinforcement learning (Barbera et al., 2016; Chen et al., 2021; Jin et al., 2014; Markowitz et al., 2018; Nonomura et al., 2018; Shin et al., 2018). We also found that there was quantitative difference in spatiotemporal organization for dSPN_ALM_ and iSPN_ALM_ (Figure 4). The correlation in dSPN_ALM_ decayed faster than iSPN_ALM_ and dSPN_ALM_ showed an ipsi-preference in the late response epoch, possibly helping to terminate contra-licks by reducing contra-preference (Figure 4C-E). Optogenetic tagging facilitates unbiased sampling of low-firing-rate neurons that allowed us to study populations of quiet neurons. The fraction of quiet neurons in the direct pathway was much higher than that in the indirect pathway. In contrast, the fraction of sensory-related neurons in the indirect pathway was larger (Figure S4). As these neurons received frontal input, the larger involvement in sensory encoding suggests that information integration in striatum had pathway preference.

Why are direct and indirect pathway SPNs co-active since they have antagonistic roles? One explanation is that the direct and indirect pathways need to function in concert to shape cortical activity to facilitate movement and cognition. Stimulation of direct and indirect pathway SPNs in anesthetized mice evokes distinct blood-oxygen-level-dependent (BOLD) signal that is widespread in regions including the basal ganglia and cortex (Lee et al., 2016). By combining stimulation of input-defined direct or indirect pathway with Neuropixel recording of cortical population activity, we showed that activation of the direct and indirect pathway oppositely shifted cortical neural trajectories underlying STM (Figure 5O, P), and the magnitude of shift predicted the degree of behavioral deficits (Figure 5T). The specific modulation of cortical population activity is not because stimulation uniformly affected cortical activity. In fact, the effects on individual cortical neurons are heterogeneous, with similar fractions of neurons up- or down-modulated. It is the specific effects on contra- and ipsi-trials in different response-types of neurons that led to the distinct modulation of neural trajectories underlying STM (Figure 7). Striatal dSPNs and iSPNs are thought to exert push-pull control over SNr by either reducing or increasing its activity. We further demonstrated that stimulation of dSPN_ALM_ and iSPN_ALM_ affected the non-selective ramping activity in SNr (Figure 6). It might seem strange that how modulation of SNr ramping activity could selectively affect cortical activity. Selective activity in ALM depends on thalamocortical projections with the ventral medial nucleus of the thalamus (VM) which receives strong GABAergic innervation from SNr (Guo et al., 2017). Non-selective activation or inactivation of SNr to VM projections specifically modulate ALM trajectories in the direction consistent with stimulation of the indirect and direct pathway, respectively (Wang et al. 2021). Modeling suggests that discrete attractor dynamics in ALM underlie STM, with an external input help to push attractors to discrete endpoints predicting different choices (Inagaki et al., 2019). The basal ganglia can function through the thalamus as an external input to modulate ALM activity to form discrete attractors.

## Acknowledgements

We thank the animal core facility at Tsinghua University for maintaining the mouse lines. This work was supported by the National Natural Science Foundation of China (32021002, 32170998).

## Author contributions

YJT and ZVG designed the project. YJT performed surgery, optogenetics and electrophysiology experiments. HJY and XC helped with mouse training, electrophysiology experiments and spike sorting. HJY performed brain transparency experiments and electrode tracks reconstruction. ZZZ and XY performed light-sheet imaging. YJT and HJY analyzed the data. XXY helped with the data analysis. YJT and ZVG wrote the paper with comments from other authors.

## Author information

The authors declare no competing interests. Correspondence and requests for materials should be addressed to.

## Materials and methods

### Animals

Mice were housed in the Animal Research Center of Tsinghua University in a 12:12 reverse light-dark cycle. All Procedures were in accordance with protocols approved by the Institutional Animal Care and Use committee at Tsinghua University, Beijing, China. This study is based on data from 47 adult mice. Three D1-cre mice (MMRRC B6.FVB(Cg)-Tg(Drd1a-cre)EY262Gsat/Mmucd) and three D2-cre mice (MMRRC B6.FVB(Cg)-Tg(Drd2-cre)ER44Gsat/Mmucd) were used to reveal spatial distribution of ALM-input defined striatal SPNs (Gong et al., 2007). Twenty D1-cre x Rosa26-LSL-FSF-ReaChR-mCit mice (JAX Stock #024846) (Hooks et al., 2015; Lin et al., 2013) and twenty-one D2-cre crossed to Rosa26-LSL-FSF-ReaChR-mCit transgenic mice were used for optogenetic stimulation of dSPN_ALM_ and iSPN_ALM_, optogenetic tagging of dSPN_ALM_ and iSPN_ALM_, SNr recording during optogenetic stimulation of dSPN_ALM_ and iSPN_ALM_, and ALM recording during optogenetic stimulation of dSPN_ALM_ and iSPN_ALM_.

### Viral injection and surgery

Adult mice were used for surgery (age > 2 months). All surgeries were done under isoflurane anesthesia (1-2%). A heat blanket was maintained at 37°C during the surgery and subcutaneous injection of Flunixin meglumine (Sichuan Dingjian Animal Medicine Co., Ltd, 1.25 mg/kg) was conducted after the surgery for 3 consecutive days to reduce inflammation.

The scalp covering the dorsal cortex of the mouse was removed. After clearing the exposed cranium, a small hole (~0.5 mm) over the left ALM (AP 2.5, ML −1.5) was made using an electric drill. Virus was injected through the small hole using a volumetric injection system (modified from Mo-10 Narishige). Glass pipettes were pulled and beveled to a sharp tip (outer diameter around 30 um), back-filled with mineral oil and front-loaded with viral suspension immediately before injection. scAAV2/1-hSyn-flex-EGFP-WPRE-pA virus (9.68E+12v.g./ml, 60nl, Taitool, Shanghai) was injected at the rate of 10 nl/min in the left ALM of D1-cre or D2-cre mice (DV 0.8) for visualization of ALM-input defined striatal direct and indirect pathways. For labelling of dSPN_ALM_ and iSPN_ALM_ with red-shifted channelrhodopsin, scAAV2/1-hSyn-Flpo-Pa virus (1.13E+13v.g./ml, 60nl, Taitool, Shanghai) was injected in the left ALM in D1-Cre x Rosa26-LSL-FSF-ReaChR-mCit mice and D2-Cre x Rosa26-LSL-FSF-ReaChR-mCit mice. Optical fibers were implanted bilaterally in SNr (AP −3.1, ML ±1.4, DV 4.2) or GPe (AP −0.4, ML ±2.0, DV 3.5) for axonal stimulation of dSPN_ALM_ and iSPN_ALM_, respectively. For optical tagging of dSPN_ALM_ and iSPN_ALM_, optrodes were advanced into the ALM-projection region of the striatum to search for light-activated units. For recording of SNr neurons during stimulation of dSPN_ALM_, optrodes were used to both activate dSPN_ALM_ axons and record SNr units.

After viral injection and fiber implantation, the bregma and craniotomy region were marked for future neurophysiology experiments. The skull was then covered with a thin layer of cyanoacrylate adhesive (Krazy glue, Elmer’s Products Inc). A titanium head-post was fixed to the posterior region and two silver pins (Digi-Key Part Number, ED90488-ND) were implanted into the cerebellum as ground and reference in electrophysiology. Dental cement was applied to fix the head-post in place.

### Behavior

The training procedure was similar as before (Guo et al., 2014a). Mice had free access to food and water for at least three days after surgery. Then mice were on food restriction for about one week before behavioral training started. The tactile-based decision task included three periods: the sample period for mice to sense whisker stimulation (1s), the delay period during which mice were trained to withhold licking (1s), and the response period for mice to lick for reward (1s). Tactile stimulus (a pole vibrated at 10Hz) was applied to the right whiskers during the sample period. The pole (about 1.2 mm in diameter) was placed ~6 mm away from the root of whiskers. Different strength of whisker stimulus was generated with different vibration amplitudes (weak: ~0.70mm; strong: ~3.16mm). After the delay period and following an auditory ‘go’ cue (0.1 s), mice licked one of the two lickports (left/right). Correct licking led to a milk reward (~4μL). Licking before the go cue (lick-early trials) during the sample or delay period restarted the period in addition to a short timeout (1s). Incorrect licking or no licking led to no reward. Mice typically had one training session (600~800 trials, ~1.5h) each day, and reached performance over 75% after 3-5 weeks’ training.

### Optogenetic stimulation of dSPN_ALM_ or iSPN_ALM_

As dSPN_ALM_ and iSPN_ALM_ spanned 2-3 millimeters in the striatum, we stimulated the projection axons in SNr and GPe, respectively. Stimulation was deployed in expert mice during sample, delay or response epoch randomly on ~25% of trials. Only one stimulation session (around 600 trials) was performed each day. Orange light from a 594 nm laser (Obis LS, Coherent) was controlled by an acousto-optic modulator (AOM; MTS110-A3-VIS, Quanta Tech) to produce a train of pulses.

The laser power was chosen not to evoke involuntary licks (5-20 mW peak power for stimulation of dSPN_ALM_ axons, 2.5-10 mW for stimulation of iSPN_ALM_ axons, 10 Hz pulse train for 1 s with each pulse lasting 2-5 ms). ‘Masking flash’ (40 × 1 ms pulses at 10 Hz, wavelength 590 nm, Luxeon Star) was applied during the whole trial period to prevent mice distinguishing control and stimulation trials (Guo et al., 2014b).

### Optical tagging of striatal dSPN_ALM_ and iSPN_ALM_

For experiments of optical tagging, D1-Cre x Rosa26-LSL-FSF-ReaChR-mCit or D2-Cre; Rosa26-LSL-FSF-ReaChR-mCit mice were injected with anterograde transynaptic virus scAAV2/1-hSyn-Flpo-Pa in the left ALM. A 64-channel silicon probes (4 shanks spaced by 250 μm, recording sites covering ~200 μm along depth, Cambridge NeuroTech) with two 100-μm-diameter fibers attached closely (<200 μm) to shanks was used for both recording and stimulation. Recordings were performed in the left hemisphere with electrodes sampling from the posterior to anterior striatum across 3 - 4 days. Craniotomy (in square shape, relative to Bregma AP: −1.0 ~ 1.5; ML: −3.5 ~ −1.0) was made two days before recording and dura of the exposed area was removed. Artificial dura and silicone elastomer (Kwik-Sil, World Precision Instrument) was used to protect the exposed area (Guo et al., 2014b). The optrode was inserted into the striatum from a chosen cortical location (e.g. AP 0.0) and moved down by at least 200 μm per recording session. At the end of the last recording session, a brief electric pulse (30 μA for 1 s, 3-4 times) was delivered to make a small lesion near the tip of the probe to facilitate reconstruction of electrode tracks (Huo et al., 2020). Each session included around 100 behavioral trials followed by optogenetic tagging using 594 nm laser light (0.5-1 mW, ~200 pulses at 2 Hz, 10 ms duration for each pulse). Higher laser powers should be avoided as strong stimulation of SPNs could inhibit recorded single-units and thus decrease the efficiency of identifying tagged dSPN_ALM_ and iSPN_ALM_ (Figure 2F).

We selected putative dSPN_ALM_ and iSPN_ALM_ based on similar criteria as in (Shin et al., 2018). First, the latency of the laser-evoked spikes was smaller than 6 ms and the latency was significantly lower (log-rank test, P < 0.01) than that without laser stimulation. Second, correlation in spike waveforms between evoked and spontaneous spikes was higher than 0.9. Third, the average activity during the tagging period (10 ms pulse) was higher than the spontaneous activity. Fourth, the peak-to-trough duration was longer than 0.6 ms to remove narrow spiking neurons that were likely fast-spiking interneurons (Figure S4A, B). In Figure 4, we used slightly relaxed criteria (P < 0.05, corr > 0.85) to reveal the spatiotemporal pattern as the results were similar (Figure S5).

### Electrophysiology

We used 64-channel silicon electrodes (Cambridge NeuroTech) and Neuropixel probes (Jun et al. 2018) for recordings in striatum, ALM and SNr. For recordings using 64-channel silicon electrodes, signals were collected by OpenEphys (http://www.open-ephys.org) or SpikeGL (Janelia Research Campus) at the sampling rate of 25 kHz. Spikes were sorted using JRCLUST (Jun et al. 2017). For recordings using Neuropixel probes, signals were collected by SpikeGLX at the sampling rate of 30 kHz and spikes were sorted by Kilosort2 (https://github.com/MouseLand/Kilosort).

Craniotomy was performed two days before recording. For ALM recording, electrodes were inserted to the left hemisphere at the angle of 14° from the vertical direction. ALM location was determined by referencing to the bregma and the maker drawn during surgery. One or two recording sessions with ~300 trials were performed each day for each mouse. Stimulation was done in the sample or delay period randomly with 25% stimulation probability. After recording in consecutive two days, mice were typically allowed to rest for one day with no recording session arranged. In total, 4-8 recording sessions were performed for each mouse. After recording, the exposed area was covered with artificial dura and silicone elastomer and Flunixin meglumine solution was subcutaneously injected.

For recordings in SNr using silicon probes, CM-DII (Invitrogen, #C7001, dissolved in absolute ethanol, 1 mg/ml) was painted on the back side of electrodes to allow visualization of electrode tracks. To record SNr neurons during stimulation of the direct pathway, optrodes were advanced into SNr to serve both stimulation and recording. Electrodes were inserted at the angle of 45° from the left hemisphere (insertion location from the cortex: AP −2.8 ~ −3.5; ML −4.0 ~ −4.6). Similar to optical tagging in striatum, electrodes were inserted to a proper place first (e.g. AP −3.3; ML −4.3; DV 2.6) and moved down by at least 200 μm after each recording session. On the second day of recording, electrodes were inserted to a more anterior region to facilitate disentangling of different recording tracks. For recordings in SNr using Neuropixel probes, electrodes were inserted progressively from the posterior to anterior or from lateral to medial parts of SNr to facilitate the separation of individual tracks.

### Electrode track reconstruction

The procedure of reconstruction of electrode tracks was similar as before (Wang et al., 2021). Basic steps included clearing of brain tissue, whole-brain imaging, and 3D reconstruction and registration to the common coordinate framework (CCF) (Wang et al., 2020).

Mouse brains were cleared following the uDISCO protocol (Pan et al., 2016). Briefly, after perfusion, mouse brains were fixed with 4% paraformaldehyde (PFA, Sigma-Aldrich Inc., St. Louis, US) in PBS (pH 7.4) at 4°C overnight. After fixation, each brain was dehydrated with serial incubations in 10-15 ml of 30 vol%, 50 vol%, 70vol%, 80 vol%, 90 vol%, 96 vol% and 100 vol% tert-butanol at 37°C water bath. Brains were immersed in 30 ml of DCM for 2 hours at room temperature for delipidation, followed by refractive index matching by immersion in 15 ml BABB-D4 solution (benzyl alcohol-benzyl benzoate and diphenyl ether mixed at a volume ratio of 4:1) at 37°C water bath for 12 h.

Cleared brains were imaged with a multi-scale light-sheet fluorescence microscope at 3 × 3 × 8 μm^3^ resolution (Zhang et al., 2021). Silicon probes and Neuropxiel probes were painted with a thin layer of CM-DiI (Invitrogen, dissolved in Ethanol, Beijing chemical works) to label recording tracks. For silicon probes, a small current (20 μA, 1 s, 4-6 times) was further delivered to produce a small lesion near the tip of the probe. And for Neuropixel probes, no lesions were made as the Neuropixel probes lack the electronic elements to pass large enough currents. Brains were imaged in two colors. Images from the blue channel (488 nm excitation) were used to mark CM-DiI tracks (and lesion sites if using silicon probes). Images from the red channel (640 nm excitation) had clear contours of brain structures and were used to register the imaged brain to the common coordinate framework (CCF, (Wang et al., 2020)). For recording using Neuropixel probes, information including LFP features, the firing rates and spike widths of SNr units was used to determine the locations of the probe tip, the surface of cortex and boundaries of specific brain regions along electrode tracks (Figure S8) (Liu et al., 2021).

For registration to CCF, the procedure was the same as before (Zhang et al., 2021). Basically, stacks of images were first stitched together to form a 3D reconstruction of the mouse brain. This process was performed automatically using Fiji plugin Grid/collection stitching ran in Matlab environment via MIJ. After stitching, the mouse brain was registered to the mouse brain template CCF (v3, http://atlas.brain-map.org/) in following steps. First, images were down-sampled at a voxel resolution of 10 × 10 × 10 μm^3^ to roughly match the size of the template brain. Second, an affine transformation was performed to correct rigid displacement and global non-rigid deformation such as stretch and shear (Advanced Normalization Tools, ANTs. Third, a b-spline transformation was performed to correct non-homogenous deformation. Forth, the combined transformation field was applied to recording tracks and lesion sites (in the case using silicon probes) to map their corresponding positions in CCF.

### Behavior data analysis

Lick-early rate was the fraction of trials that mice licked before the response period. No-response rate was the fraction of trials that mice did not lick during the response period. Licking frequency was the number of licks per unit of time (Figure 1J, binsize, 1 ms; smooth window, 200 ms). Performance was calculated as the fraction of correct trials, excluding the lick-early and no-response trials. Sessions with control performance less than 70% were excluded. For performance during stimulation in the left or right hemisphere (Figure 1H, Figure S3B), ipsi and contra denoted directions relative to the recorded hemisphere. Performance bias caused by stimulation was calculated as performance in contra-trials subtracted by performance in ipsi-trials (Figure 5T, Figure S7N).

### Electrophysiological data analysis

Neurons were tested for trial-type selectivity by comparing firing rates in ipsi- and contra-trials during sample, delay and response periods. Neurons that significantly differentiated trial-types during sample, delay or response epoch were defined to be sample-selective, delay-selective or response-selective (selected using t-test with P < 0.05). Selective neurons were further classified to be ipsi-preferring and contra-preferring if total spike counts in the specified period were higher in ipsi-trials and contra-trials, respectively. Preferred-trials were trials during which ipsi-preferring or contra-preferring neurons had higher spike counts while during non-preferred trials ipsi-preferring or contra-preferring neurons had lower spike counts. Selectivity was defined as the difference of spike counts between preferred and non-preferred trials (Figure 5J, M; Figure 6I, L). The averaged selectivity was the mean absolute firing rate difference between the preferred and non-preferred trials (Figure 3F). To compute ‘normalized contra-selectivity’, we normalized the selectivity by its peak value (Figure 3C).

The coding direction (CD) is a n dimensional vector in activity space that maximally distinguish contra-trials and ipsi-trials (Li et al., 2016). Here, n is the number of simultaneously recorded neurons and we selected sessions with n ≥ 5 as smaller simultaneously recorded neurons will make dimensionality reduction unstable. For each session, we randomly selected half of trials to compute CD at each time point (20 ms bin size) and projected the remaining half of trials to the CD to obtain the projected trajectories (Figure 5, 6). The neural trajectories were smoothed with a 400 ms time window. To obtain the ramping mode (RM), we first combined the neural activity from both trial types to form the population activity matrix (n x trial-number). Then we calculated the eigenvectors of the population activity matrix using Singular Value Decomposition (SVD). The eigenvectors were rotated using the Gram-Schmidt process to be orthogonal to CD and to each other. The projection to the first vector resulted in nonselective ramping activity, and this vector was referred to as the ramping mode (Inagaki et al., 2018). The RM-projected trajectories were also smoothed with a 400 ms time window. To examine the correlation of trajectory shift and performance change, we calculated the CD change as the difference between projected trajectories in control and stimulation trials (average during the last 300 ms of the delay period). The CD change in ipsi- and contra-trials were averaged. The positive (or negative) shift represents the change of trajectory along ipsi- to contra-(or contra- to ipsi-) direction (Figure 5T). The decision boundary (DB) was defined as:

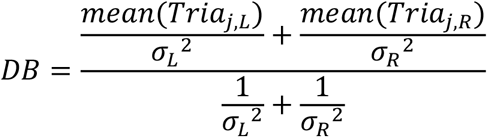

where *mean*(*Tria*_*j*_) represents the mean projected value of each trial on CD. *L*, *R* represent trial types and *σ*^2^ is the variance of projected value. For each session, we calculated the DB based on half of the trials and obtained the distribution of distance between projected value and DB for the other half of trials (Figure 5U).

To reveal the spatiotemporal distribution of optogenetically identified SPNs, selectivity within each time window was normalized to the maximum selectivity at the selected plane (Figure 4E, F). Neurons with positive selectivity were contra-selective neurons while those with negative selectivity represented ipsi-selective neurons. Summed selectivity ratio was the summed ipsi-selectivity divided by summed contra-selectivity of all neurons.

---

**Figure S1.**
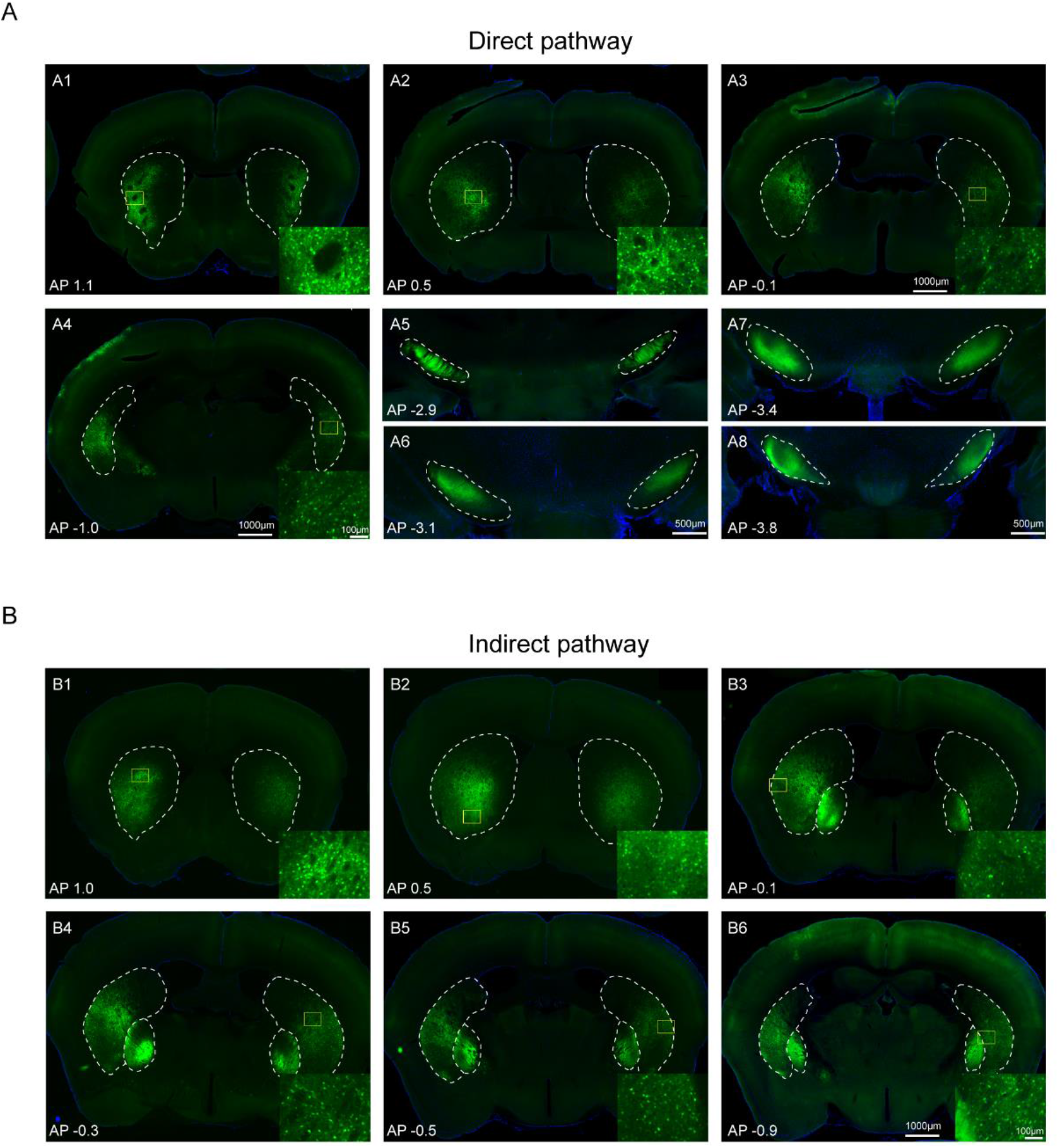
Widespread expression of dSPN_ALM_ and iSPN_ALM_ neurons. (A) Striatal dSPN_ALM_ neurons and their axons in the ipsilateral and contralateral hemispheres. (B) Striatal iSPN_ALM_ neurons and their axons in the ipsilateral and contralateral hemispheres. Scale bars in A1-A4 and B1-B6, 1000 μm. Scale bars in A5-A8, 500 μm. Scale bars in inset, 100 μm.

**Figure S2.**
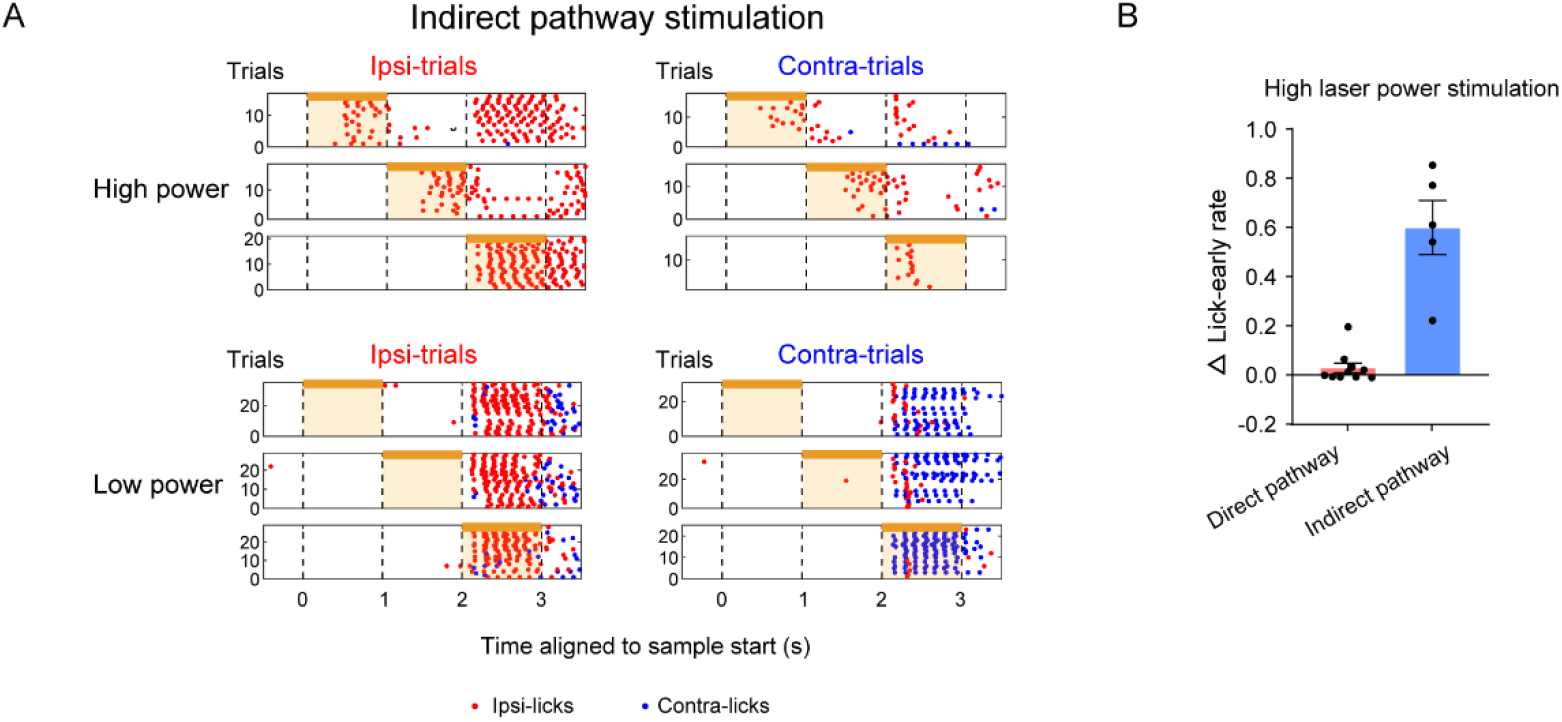
High-power stimulation of the indirect pathway induces early licks. (A) Licking pattern of an example mouse during indirect pathway stimulation with high (top) and low (bottom) laser power, respectively (high laser power, a 40Hz pulse train with 20 mW peak power; low laser power, a 10Hz pulse train with 5mW peak power; pulse duration, 2ms). (B) Lick-early rate during stimulation of direct and indirect pathways with high laser power. Each dot represents a behavioral session. Laser power, 20 mW peak power for direct pathway stimulation; 3-20 mW peak power for indirect pathway stimulation (the peak power was variable as relatively lower laser powers could evok early licks in some mice).

**Figure S3.**
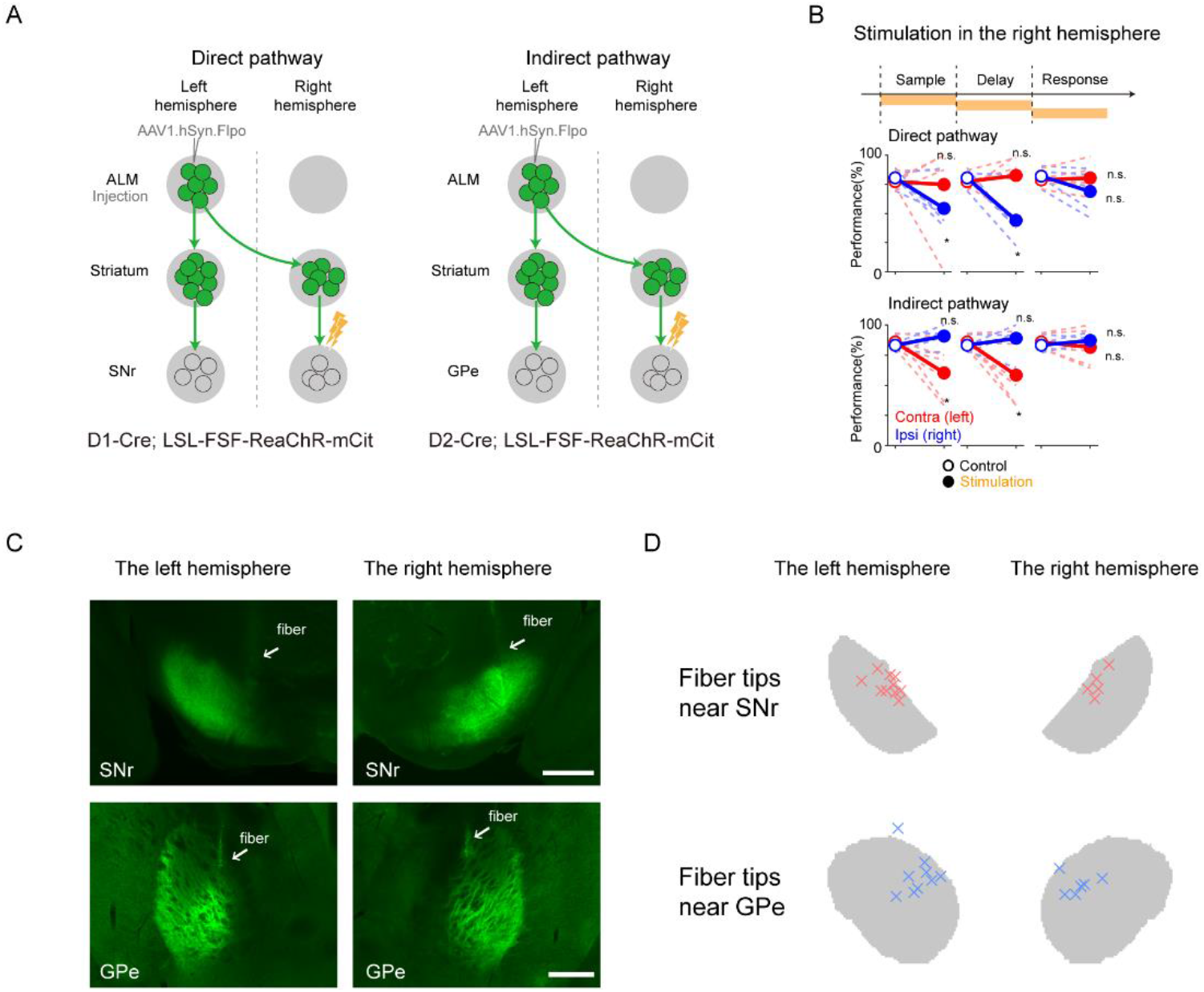
Stimulation of ALM-input defined direct and indirect pathway SPNs in the right hemisphere. (A) Schematic of stimulation of axons of dSPN_ALM_ (left) and iSPN_ALM_ (right) in the right hemisphere (contralateral to the side of viral injection). (B) Stimulation of dSPN_ALM_ in the right hemisphere produced a contralateral bias, while stimulation of iSPN_ALM_ in the right hemisphere produced an ipsilateral bias (contra and ipsi denote the side relative to the stimulated right hemisphere). Thus stimulation of ALM-input defined direct and indirect pathways produced opposite directional bias. The direction of bias (relative to the stimulated hemisphere) is the same between stimulations in the left and right hemispheres. Thin line, individual mice (n = 5 D1-Cre, n= 6 D2-Cre mice). Thick line, mean. (C) Axonal projections of dSPN_ALM_ (left) and iSPN_ALM_ (right) in the left and right hemispheres. Fiber locations were determined based on autofluorescence of fiber tracks near SNr and GPe. (D) Compilation of fiber tip positions for stimulation of the direct and indirect pathways (n = 10 for the left SNr; n = 5 for the right SNr; n = 9 for the left GPe; n=6 for the right GPe). Scale bar, 500 μm. * denote paired two-sided t-test with statistical significance P < 0.05. ns, not significant.

**Figure S4.**
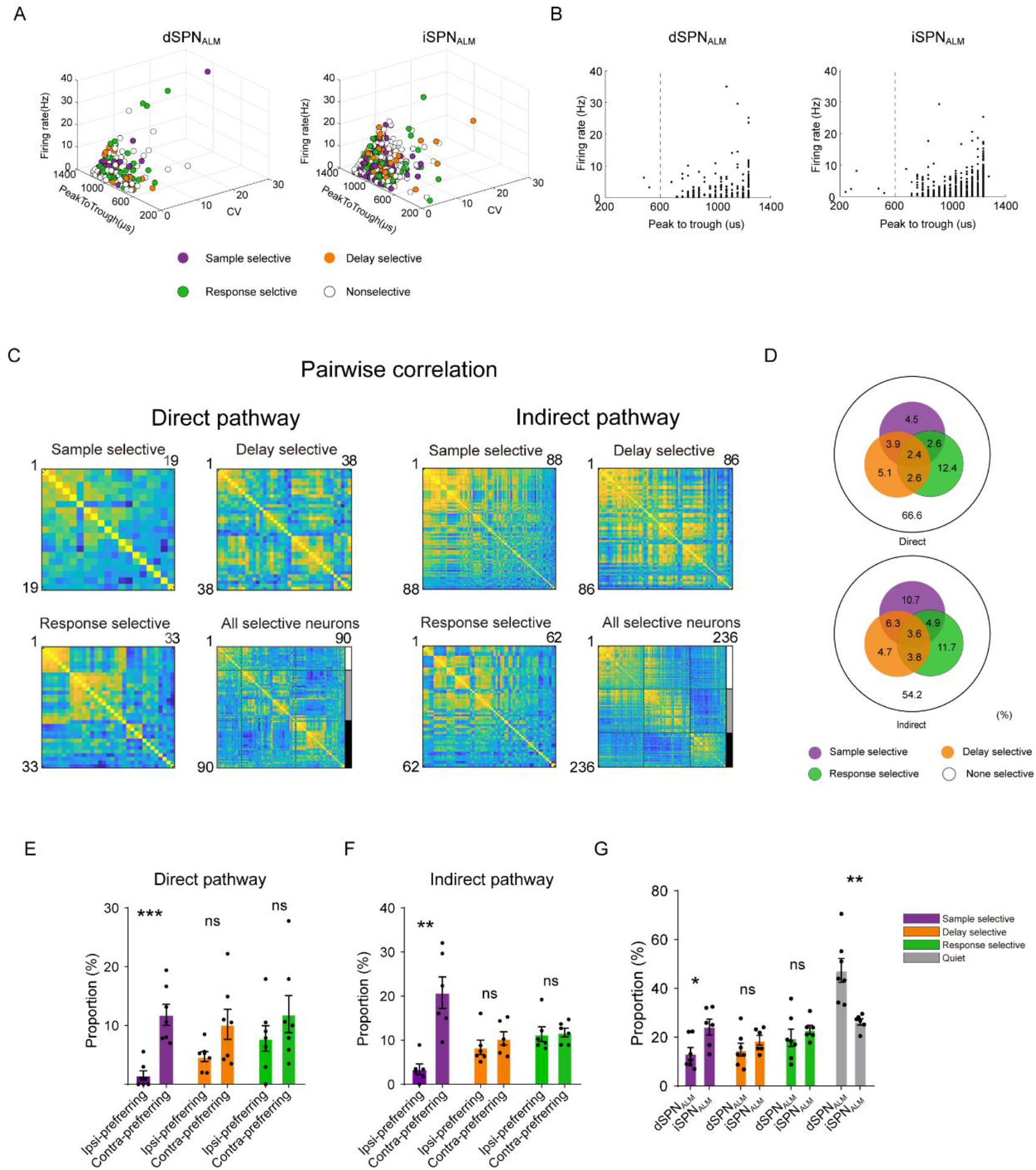
Property of optogenetically identified dSPN_ALM_ and iSPN_ALM_. (A) Neurophysiological properties of identified dSPN_ALM_ (left) and iSPN_ALM_ (right). CV, coefficient of variation of spike intervals. Sample-, delay- or response-selective neurons are indicated with distinct colors. (B) Scatter plot of firing rates and peak-to-trough durations for dSPN_ALM_ (left) and iSPN_ALM_ (right). (C) Pairwise correlation of sample-selective, delay-selective or response-selective neurons in the direct and indirect pathways. (D) Percentage of different response-types of dSPN_ALM_ (top) and iSPN_ALM_ (bottom). (E) Proportion of ipsi- and contra-preferring neurons of the direct pathway. Purple, sample selective; orange, delay selective; green, response selective. There were more contra-preferring sample-selective neurons. (F) Similar to (D) but for the indirect pathway. There were more contra-preferring sample-selective neurons. (G) Comparison of sample-, delay- and response-selective neurons, and quiet neurons in the direct and indirect pathway. *, **, & *** denote two-sided t-test with statistical significance P < 0.05, P < 0.01, & P < 0.001, respectively. ns, not significant.

**Figure S5.**
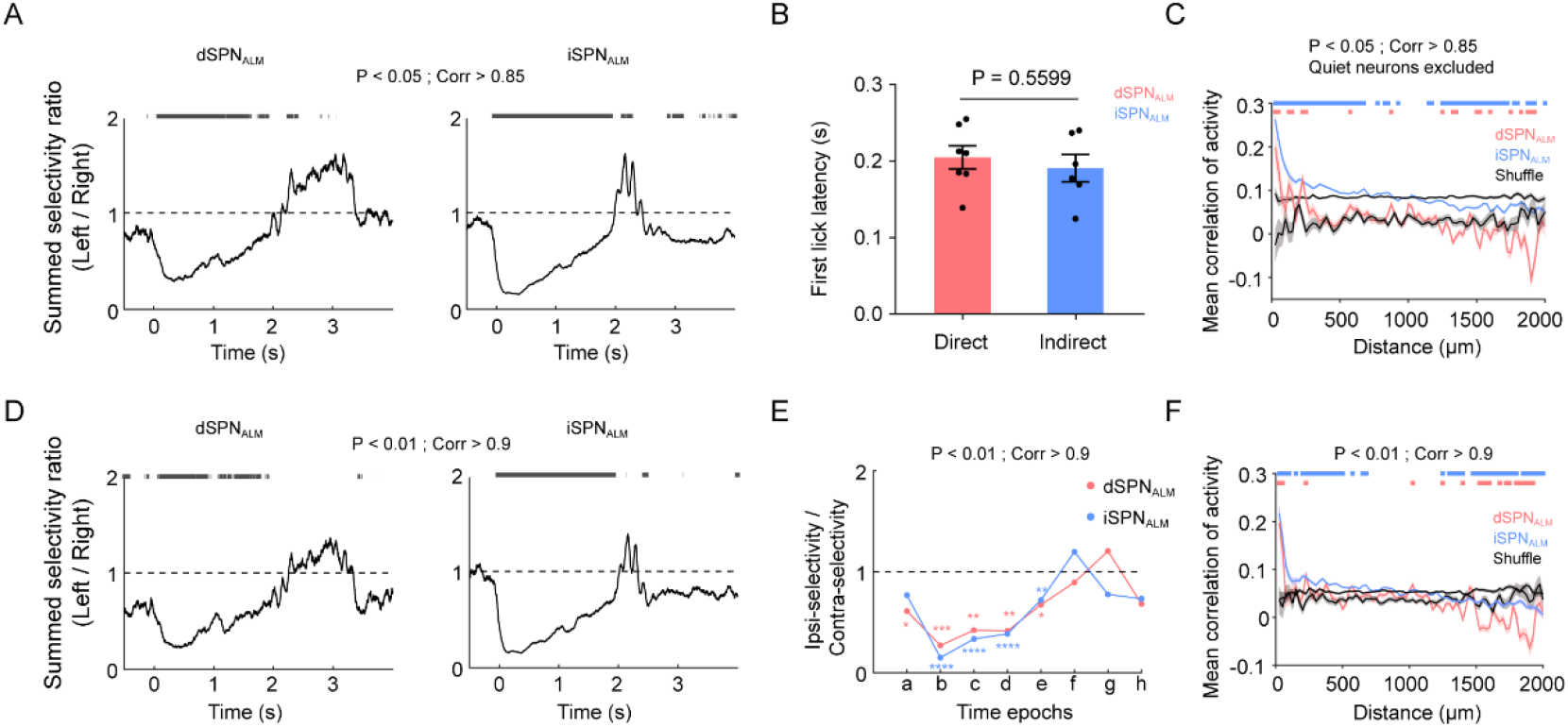
Selectivity ratio of direct and indirect pathway SPNs. (A) Selectivity ratio of dSPN_ALM_ (left) and iSPN_ALM_ (right). The black bar indicates duration of significant difference from 1 (two-sided t-test, P < 0.05). Smooth window, 200 ms. Criteria in selecting tagged neurons, P < 0.05 and Corr >0.85. (B) First lick latency in D1-Cre and D2-Cre mice used for optogenetic tagging of dSPN_ALM_ and iSPN_ALM_. Each dot represents one mouse. (C) Correlation of activity in dSPN_ALM_ (light red) and iSPN_ALM_ (light blue). Same format as in Figure 4C but with quiet neurons removed. (D) Selectivity ratio of dSPN_ALM_ (left) and iSPN_ALM_ (right). Same format as in (A) but with more restricted criteria in selecting tagged neurons (P < 0.01, Corr >0.9). (E) Summed selectivity ratio of dSPN_ALM_ and iSPN_ALM_ in eight time epochs. Same format as in Figure 4D. Same Data as in (D). (F) Correlation of activity in dSPN_ALM_ (light red) and iSPN_ALM_ (light blue). Same format as in Figure 4C. Same Data as in (D). *, **, ***& **** denote two-sided t-test with statistical significance P < 0.05, P < 0.01, P < 0.001 & P < 0.0001, respectively.

**Figure S6.**
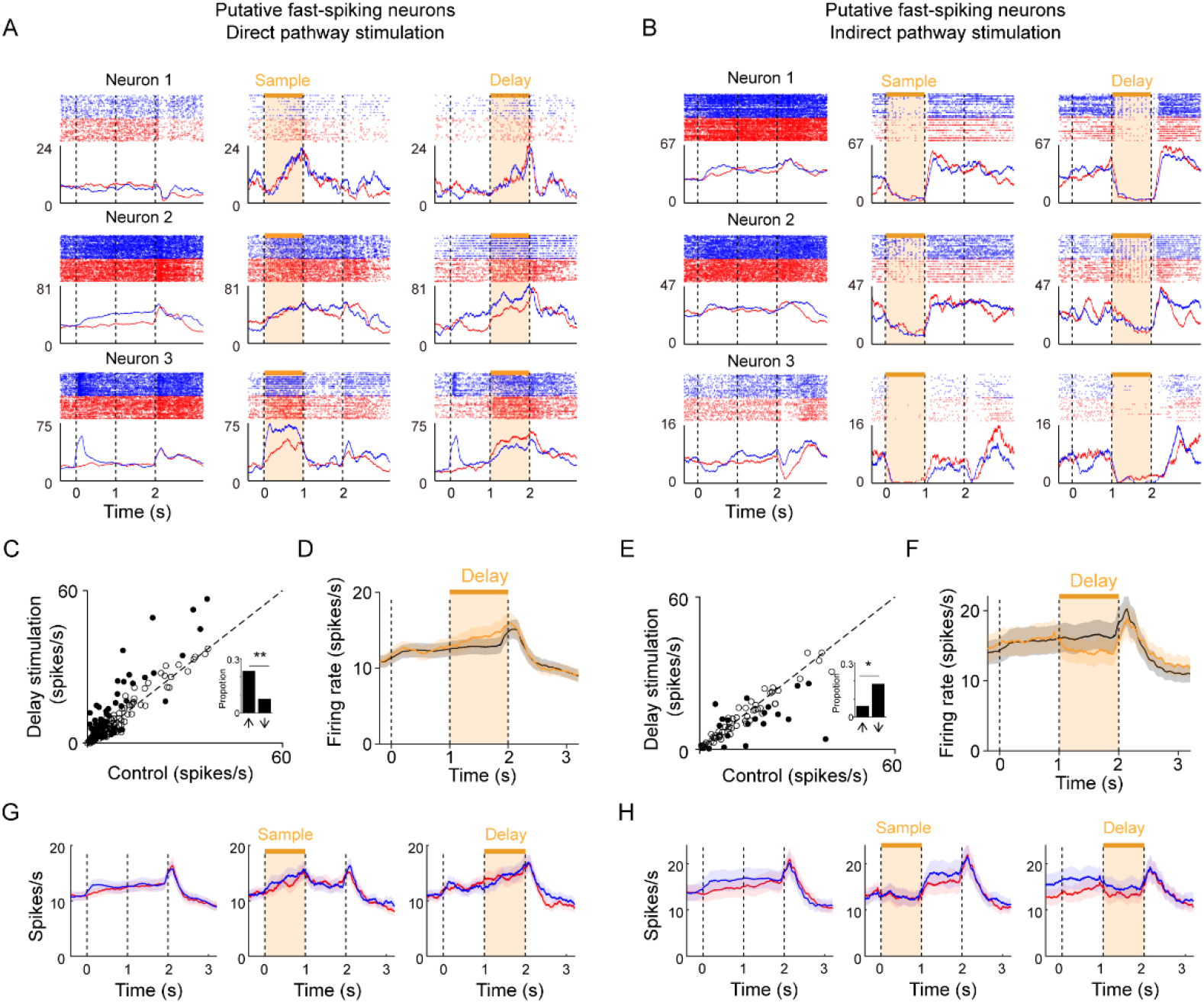
Stimulation of ALM-input defined direct and indirect pathways oppositely but less selectively modulate fast spiking neurons (FSN) in ALM. (A) Activity of three example FSNs during stimulation of the direct pathway. Each row shows one FSN. Left, control trials. Middle, trials with stimulation during the sample period. Right, delay-period stimulation. (B) Same format as in (A) but for FSNs during stimulation of the indirect pathway. (C) Scatter plot of mean firing rates of putative FSNs in ALM (119 units from 7 mice). Stimulation of the direct pathway during the delay epoch mainly increased rather than decreased FSN activity. Filled circles, neurons that were significantly modulated (selected using t-test with significance P < 0.05). Dotted line is the unity line. Inset, proportion of up-modulated and down-modulated neurons. (D) Average activity of FSNs in control (black) and stimulation (orange) conditions. Mean activity were significantly increased (by 11.2 ± 4.9%, P < 0.05, paired t-test). (E) Stimulation of the indirect pathway during the delay epoch mainly decreased rather than increased FSN activity (73 units from 4 mice). Same format as in (C). (F) Same format as in (D) but for FSNs during stimulation of the indirect pathway. Mean activity were significantly decreased (by 14.4 ± 6.3%, P < 0.05, paired t-test). (G) Average activity of FSNs in ipsi-trials (red) and contra-trials (blue) during stimulation of the direct pathway. Left, control trials. Middle, trials with stimulation during the sample period. Right, delay-period stimulation. (H) Same format as in (G) but for stimulation of the indirect pathway. * & ** denote Chi-square test with statistical significance P < 0.05 & P < 0.01, respectively.

**Figure S7.**
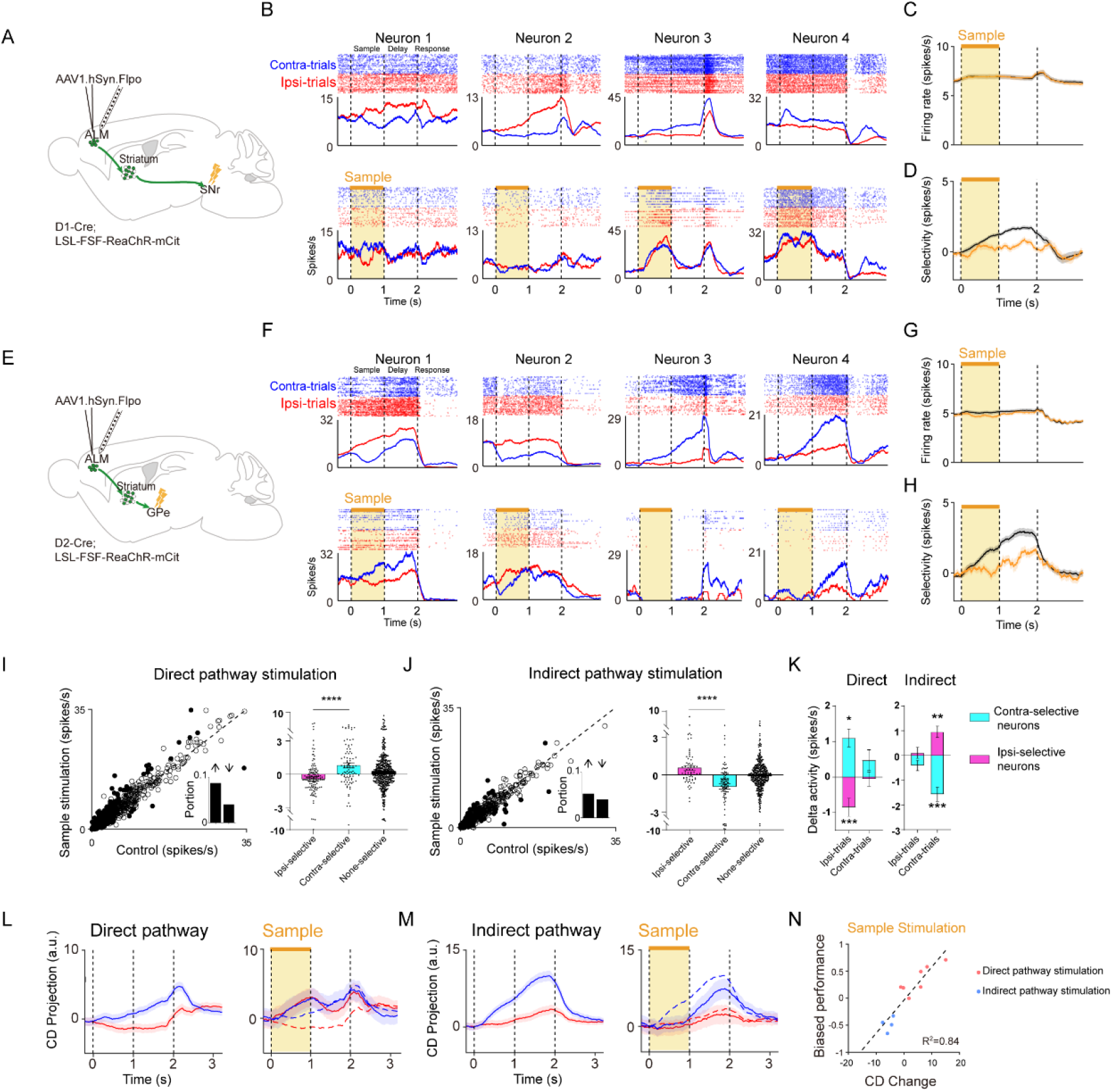
Stimulation of ALM-input defined direct and indirect pathways during the sample period oppositely modulate ALM activity. Same format as in Figure 5, but for stimulation during the sample period. For (I-K), we analyzed the effect of stimulation on activity during the delay epoch. For (R) we analyzed the correlation between behavior bias and CD change during the last 300ms of the delay period. *, **, *** & **** denote two-sided t-test with statistical significance P < 0.05, P < 0.01, P < 0.001& P < 0.0001, respectively.

**Figure S8.**
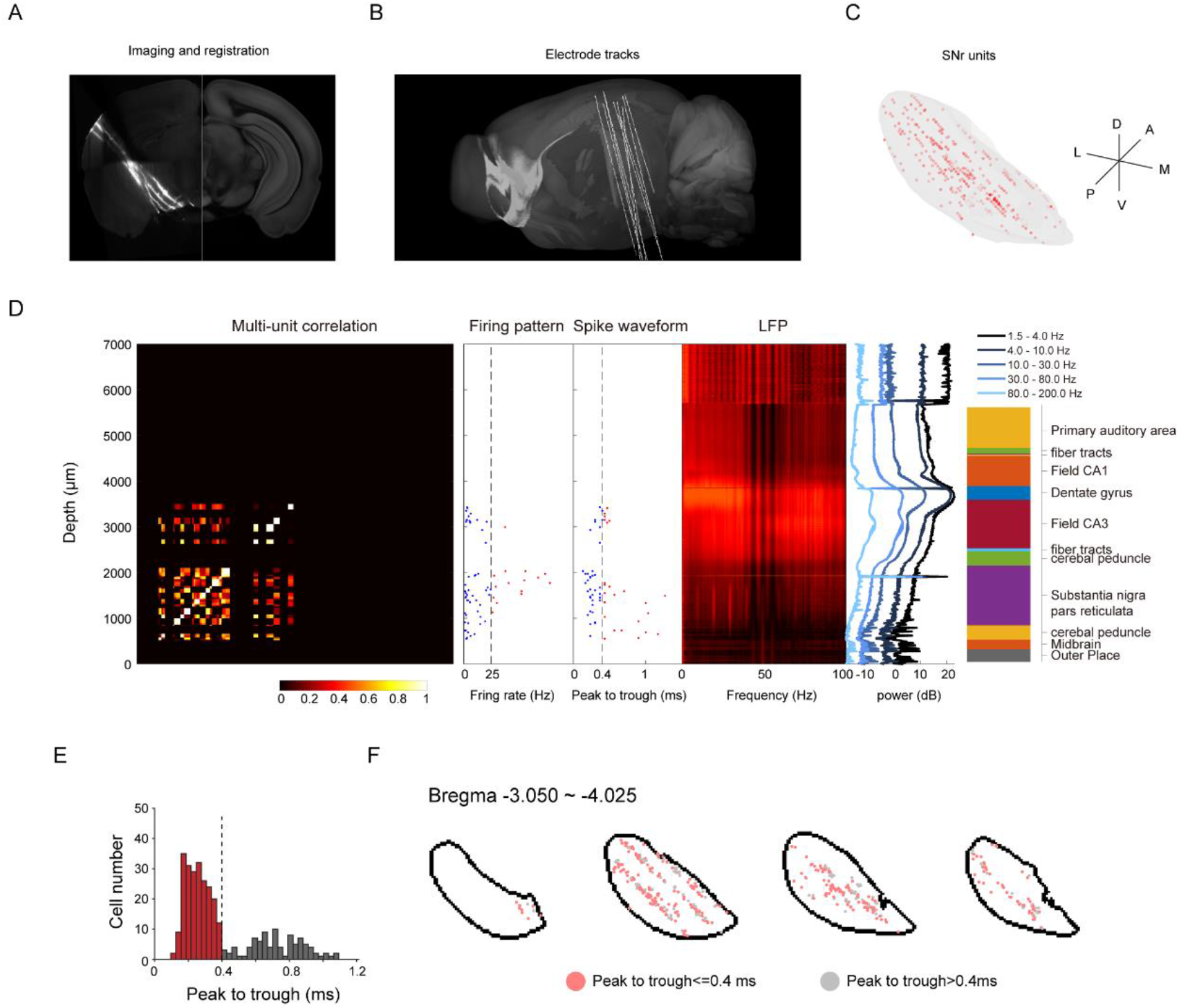
Reconstruction of recording locations in SNr for experiments using Neuropixel probes. (A) Registration of whole brain imaging data to the Allen CCF (similar to Figure 4A). (B) Example electrode tracks. (C) Distribution of recorded SNr units. Each dot represents one unit. (D) Multiple neurophysiology landmarks along Neuropixel probes were used to register electrode tracks to the Allen CCF (see Methods). Left, correlation of multi-unit activity from 384 recording sites near the probe tip (near or within SNr). Middle, firing rates and spike widths of sorted single-units. Right, LFP recording from 768 recording sites after the behavioral session and distribution of power within different frequency bands. (E) Spike widths (peak to trough) of units near SNr form bimodal distribution. Red, putative GABAergic neurons selected based on the distribution. Gray, unclassified. (F) Locations of selected SNr units projected on different coronal sections (units recorded using Neuropixel probes only).

